# Role of Forkhead box F1 in the Pathobiology of Pulmonary Arterial Hypertension

**DOI:** 10.1101/2024.09.18.611448

**Authors:** Jose Gomez-Arroyo, Arjan C. Houweling, Harm Jan Bogaard, Jurjan Aman, Joseph A. Kitzmiller, Aleksey Porollo, Dennis Dooijes, Lilian J. Meijboom, Phillip Hale, Michael W. Pauciulo, Jason Hong, Na Zhu, Carrie Welch, Yufeng Shen, William J. Zacharias, Francis X. McCormack, Micheala A. Aldred, Matthew T. Weirauch, Stefan Graf, Christopher Rhodes, Wendy K. Chung, Jeffrey A. Whitsett, Lisa J. Martin, Vladimir V. Kalinichenko, William C. Nichols

## Abstract

**Rationale:** Approximately 80% of patients with non-familial pulmonary arterial hypertension (PAH) lack identifiable pathogenic genetic variants. While most genetic studies of PAH have focused on predicted loss-of-function variants, recent approaches have identified ultra-rare missense variants associated with the disease. *FOXF1* encodes a highly conserved transcription factor, essential for angiogenesis and vasculogenesis in human and mouse lungs.

**Objectives:** We identified a rare *FOXF1* missense coding variant in two unrelated probands with PAH. *FOXF1* is an evolutionarily conserved transcription factor required for lung vascular development and vascular integrity. Our aims were to determine the frequency of *FOXF1* variants in larger PAH cohorts compared to the general population, study *FOXF1* expression in explanted lung tissue from PAH patients versus control (failed-donor) lungs, and define potential downstream targets linked to PAH development.

**Methods:** Three independent, international, multicenter cohorts were analyzed to evaluate the frequency of *FOXF1* rare variants. Various composite prediction models assessed the deleteriousness of individual variants. Bulk RNA sequencing datasets from human explanted lung tissues were compared to failed-donor controls to determine *FOXF1* expression. Bioinformatic tools identified putative *FOXF1* binding targets, which were orthogonally validated using mouse ChIP-seq datasets.

**Measurements and Main Results:** Seven novel or ultra-rare missense coding variants were identified across three patient cohorts in different regions of the *FOXF1* gene, including the DNA binding domain. *FOXF1* expression was dysregulated in PAH lungs, correlating with disease severity. Histological analysis showed heterogeneous *FOXF1* expression, with the lowest levels in phenotypically abnormal endothelial cells within complex vascular lesions in PAH samples. A hybrid bioinformatic approach identified FOXF1 downstream targets potentially involved in PAH pathogenesis, including *BMPR2*.

**Conclusions:** Large genomic and transcriptomic datasets suggest that decreased *FOXF1* expression or predicted dysfunction is associated with PAH.

## Introduction

Pulmonary arterial hypertension (PAH) is a rare lung vascular disease frequently complicated by right ventricular (RV) dysfunction and failure(1). Despite remarkable therapeutic advances, PAH remains an incurable, progressive disease which frequently leads to death within 3-10 years of diagnosis(2). The complexity of the pulmonary vascular system poses a significant challenge in identifying *bona fide* cellular and molecular pathogenetic targets responsible for disease progression. Whereas many cell types have been implicated in PAH(3–5), endothelial cell (EC) dysfunction, injury and death have emerged as key drivers in the pathogenesis of the disease(3, 6). Many conditions characterized by chronic EC injury are associated with PAH, including drug toxicity(7–11), chronic HIV infection(12), autoimmune diseases(13) and chronic interleukin-mediated inflammation(14– 17). Conversely, inhibiting EC apoptosis in a pre-clinical model of PAH prevents the development of pathological vascular remodeling(18).

The predisposing factors causing EC dysfunction, injury or death and the mechanistic link to irreversible vascular remodeling in PAH are incompletely understood. Extensive genomic analysis of families and large cohorts of patients with PAH has identified pathogenic variants in different genes that could predispose to EC dysfunction or premature EC death(19). Heterozygous germline mutations in the bone morphogenetic protein receptor II (*BMPR2*) gene are the most prevalent in the heritable form of PAH; however, based on clinical observations and experimental evidence several additional genes have been associated with heritable PAH including

## Results

### Genotype-phenotype Analysis and Pedigree of Index Cases Carrying a Rare FOXF1 Variant

The first index case (proband-1, III:1) that focused our interest on *FOXF1* was an 18-year-old female diagnosed with idiopathic PAH who presented with long-standing complaints of shortness of breath(33). Initial work up revealed elevated pulmonary artery systolic pressure by transthoracic echocardiography, and severely dilated right-sided heart chambers. Cardiac magnetic resonance imaging (MRI) demonstrated severe right ventricular (RV) dilation and hypertrophy (**Figure 1A-B**). Right heart catheterization confirmed the diagnosis of severe PAH *ACVRL1, GDF2, KCNK3, ENG, ATP13A3, CAV1, TBX4, SOX17, CAPNS1* and *KDR*(20–25). Here we present evidence supporting Forkhead box F1 (*FOXF1*) as a new risk-gene involved in the pathobiology of PAH. *FOXF1* encodes a highly conserved transcription factor, essential for neonatal angiogenesis in human and mouse lungs(26, 27). *FOXF1* regulates the expression of many key ligands and receptors required for EC function and survival(26, 28). While deletions or loss-of-function mutations in *FOXF1* are associated with fulminant pulmonary hypertension in neonatal patients with alveolar capillary dysplasia with misalignment of pulmonary veins (ACDMPV)(26, 29, 30), the role of *FOXF1* in adult PAH remains largely unknown.

**Figure 1:**
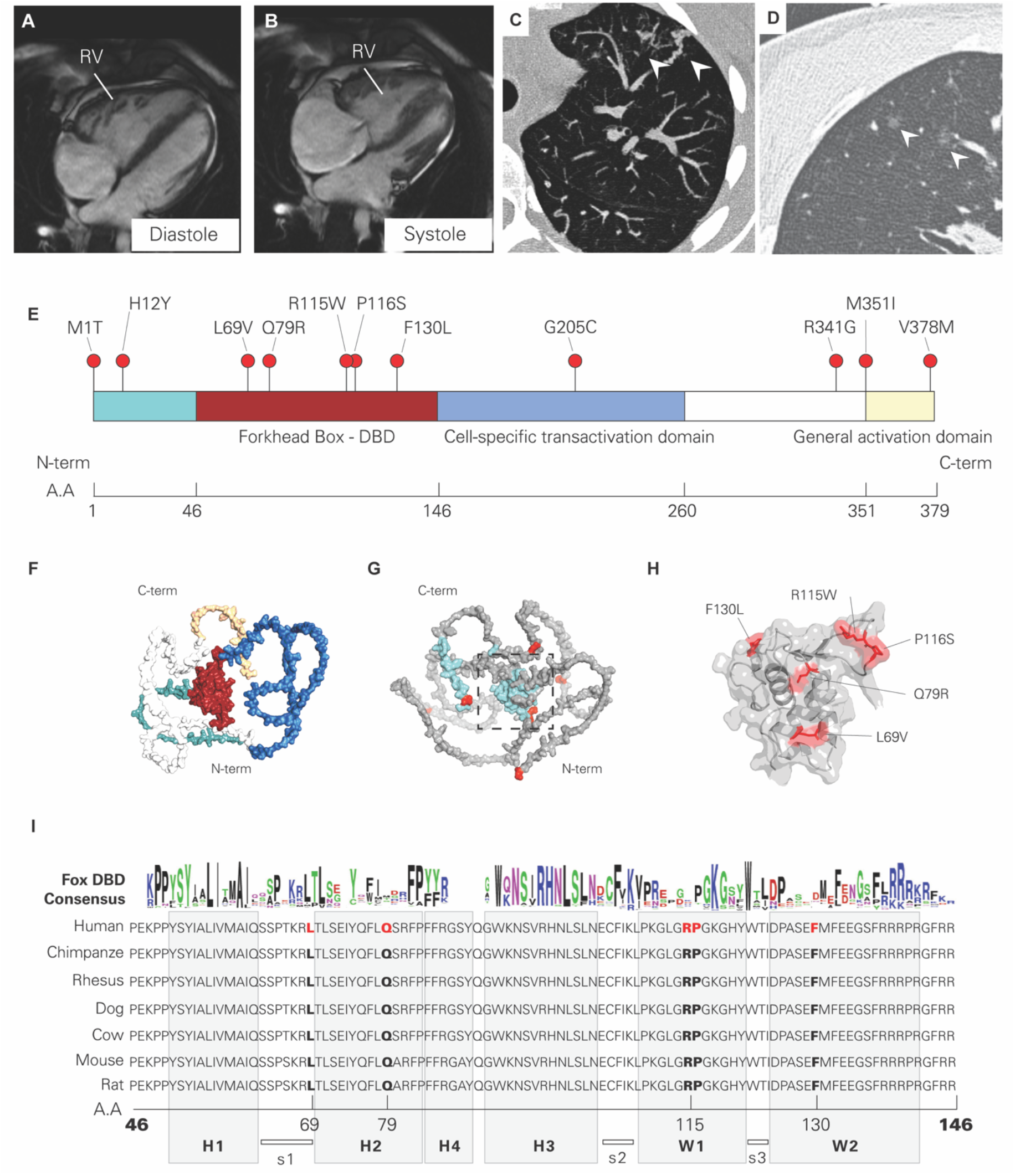
FOXF1 variants in patients with pulmonary arterial hypertension. A-B) Cardiac MRI shows a dilated and hypertrophic right ventricle (RV) during systole and diastole in a patient with PAH and severe RV failure, carrying a deleterious missense *FOXF1* variant c.1021C>G, p.R341G. C) computed tomography pulmonary angiography illustrates abnormal distal pulmonary vasculature with severe tortuous arterial vessels in the left upper lobe and in the right upper lobe centrilobular ground glass nodules (D, arrows). E) Lollipop chart indicating the location of every variant found in PAH patients and their relative sequence location along the protein domains of FOXF1. F) Three-dimensional structure of FOXF1 predicted by AlphaFold2. Different regions are colored according to the color scheme used in panel E. G) Variants found in PAH projected within the three-dimensional structure of FOXF1. Cyan represents highly conserved regions across species and red represents mutated residues in PAH patients, shown in panel E. H) Magnified view of the DNA binding domain with mutated residues shown in red. I) Amino acid sequence of the DNA binding domain with PAH-associated mutated residues in red. A sequence logo representing the residue conservation of human FOX-DBDs, using sequence alignment profile of all the 50 human FOX-DBDs, modified from Dai, et al(69)) is shown at the top to illustrate the residue conservation.

In the present study, we describe the clinical course of two adult patients diagnosed with PAH, carrying rare, non-synonymous coding variants in *FOXF1*. Additional queries identified several missense coding *FOXF1* variants in three independent, multicenter U.S. and European cohorts(20, 21, 31), including patients with idiopathic, familial, and associated (secondary) forms of PAH. *FOXF1* was identified as one of the most downregulated genes in transcriptome-wide, bulk RNA sequencing analysis from a separate U.S. tissue biobank, composed of explanted lung tissue samples derived from PAH patients and failed-donor control subjects(32). Deep phenotyping enabled us to provide statistical evidence for an association between *FOXF1* lung tissue expression and disease severity. Lastly, using a hybrid bioinformatics approach, we identified predicted *FOXF1* downstream targets with potential involvement in the pathogenesis of PAH, including *BMPR2*.

(right atrial pressure of 13 mmHg, mean pulmonary artery pressure [mPAP] of 95 mmHg, cardiac index of 3.1 l/min/m^2^, pulmonary capillary wedge pressure of 13 mmHg and a pulmonary vascular resistance of 17 Wood units). She denied any history of smoking, intravenous drug use, amphetamines, methamphetamines, or cocaine. Extensive work up was negative for comorbidities associated with PAH, and thus, a diagnosis of idiopathic PAH was made. Computed tomography pulmonary angiography demonstrated anastomoses between venous structures and bronchial arteries, as well as thickened interlobular septae and discrete centrilobular ground glass opacities (**Figure 1C-D**) concerning for pulmonary venoocclusive disease, which prompted genetic testing. A whole exome sequencing (WES)-based panel revealed heterozygosity for a rare non-synonymous coding variant in *FOXF1* (c.1021C>G, p.Arg341Gly). Secondary whole exome data analysis did not reveal additional variants of interest or pathological variants in PAH-associated genes. The proband-1’s sister (III:2), mother (II:1) and father (II:2) are alive and without apparent cardiopulmonary disease. A maternal first cousin (III:3) died at the age of 3.5 years from PAH, which was attributed to an atrial septal defect. Trio-WES confirmed the same *FOXF1* variant (c.1021C>G, p.Arg341Gly) in the proband-1’s unaffected father but not in her mother. No genotype data were available for the other family members. A diagram of the extended pedigree summarizing two generations carrying the genetic variant is shown in **Supplemental Figure 1C**.

The second index case (proband-2, III:1) involved a 25-year-old male with a history of recurrent airway infections and IgA deficiency, who was diagnosed with precapillary pulmonary hypertension via right heart catheterization (mean pulmonary artery pressure of 54 mmHg, pulmonary capillary wedge pressure of 7 mmHg, cardiac output 6.5 L/min, and a pulmonary vascular resistance of 7.2 Wood units). Pulmonary angiography demonstrated homogeneous peripheral perfusion defects, consistent with thrombotic angiopathy and distal vascular pruning, raising concerns for group I PH (PAH). Genetic testing with a WES-based PAH panel demonstrated the same rare variant in *FOXF1* (c.1021C>G, p.Arg341Gly) found in proband-1, without other variants in PAH-associated genes. His family history was negative for PAH. The proband-2’s mother (II:1), father (II:2), and two brothers (III:3 and III:4) are alive and without cardiopulmonary disease. Trio-WES demonstrated the same single nucleotide variant (SNV) in *FOXF1* in proband-2’s unaffected father but not in his mother. A pedigree chart summarizing two generations carrying the genetic variant is shown in **Supplemental Figure 1D**.

The *FOXF1*: c.1021C>G, p.R341G variant found in both probands is ultra-rare based on a global minor allele frequency (MAF) of *f*= 0.000016 and a European (Non-Finnish) MAF of *f*= 0.000035, estimated from the Genome Aggregation Database (gnomAD exomes, v2.1) (34). The very-low allele frequency coincided with two critical mutational constraint metrics used to score gene essentiality, suggesting poor mutation tolerability for *FOXF1* in the general population(34): 1) intolerance to heterozygous predicted loss-of-function variation (pLI) value of 0.96, and 2) loss-of-function observed/expected and upper bound fraction (LOEUF) of 0 [0 -0.3].. **Table 1 and Supplemental Table 3** summarize the results from different *in-silico* tools used to predict the functional impact and potential impact of the *FOXF1* variant c.1021C>G, p.Arg341Gly. Notably, the REVEL score was 0.66 and the CADD score was 23. The latter is a broadly used tool to estimate the deleteriousness of single nucleotide variants in the human genome, based on the principle that variants reducing organismal fitness are typically depleted by natural selection(35, 36). A variant with a CADD (scaled C-score) of 20 or higher indicates that these variants are predicted to be the 1% most deleterious substitutions possible in the human genome(36). A CADD score of 23 for the c.1021C>G, p.R341G *FOXF1* variant coincides with the very-low allele frequency estimated by gnomADv2.1.1, and all together, highly suggested a deleterious role in these two index cases with PAH.

**Table 1.** Summary of the algorithms predicting molecular pathogenicity for all FOXF1 missense variants included in this study.

| Proband Number | Coding Impact | Chr | Position | cDNA | Protein | Protein Domain Location | GnomAD MAF* | PolyPhen Cat | SIFT Cat | M-CAP | MetaSVM | MetaRNN | REVEL | CADD v1.7 | AlphaMissense | Alpha Missense Ca | ACMG Classification |
| --- | --- | --- | --- | --- | --- | --- | --- | --- | --- | --- | --- | --- | --- | --- | --- | --- | --- |
| 1 | Missense | 16 | 86512966 | c.1021C>G | p.(Arg341Gly) | General AD | 0.00000657 | Probably damaging | Deleterious | Pathogenic Moderate | Pathogenic Moder | Pathogenic Moderate | 0.663 | 22.4 | 0.9164 | Likely Pathogenic | VUS |
| 2 | Missense | 16 | 86512966 | c.1021C>G | p.(Arg341Gly) | General AD | 0.00000657 | Probably damaging | Deleterious | Pathogenic Moderate | Pathogenic Moder | Pathogenic Moderate | 0.663 | 22.4 | 0.9164 | Likely Pathogenic | VUS |
| 3 | Missense | 16 | 86510912 | c.343C>T | p.(Arg115Trp) | Forkhead DBD | 0 | Probably damaging | Deleterious | Pathogenic Moderate | Pathogenic Moder | Pathogenic Moderate | 0.658 | 28.6 | 0.993 | Likely Pathogenic | VUS |
| 4 | Missense | 16 | 86510959 | c.390C>G | p.(Phe130Leu) | Forkhead DBD | 0 | Possibly damaging | Deleterious | Pathogenic Supp | Uncertain | Uncertain | 0.631 | 24 | 0.9958 | Likely Pathogenic | VUS |
| 5 | Missense | 16 | 86510774 | c.205C>G | p.(Leu69Val) | Forkhead DBD | - | Possibly damaging | Tolerated | Pathogenic Supp | Pathogenic Moder | Pathogenic Strong | 0.809 | 20 | 0.7086 | Likely Pathogenic | VUS |
| 6 | Missense | 16 | 86510571 | c.2T>C | p.(Met1Thr) | N/A | 0 | Benign | Deleterious | Pathogenic Moderate | Uncertain | Pathogenic Strong | 0.509 | 22.2 | Not availabl | Not available | VUS |
| 7 | Missense | 16 | 86510915 | c.346C>T | p.(Pro116Ser) | Forkhead DBD | 0 | Probably damaging | Deleterious | Pathogenic Moderate | Pathogenic Moder | Uncertain | 0.803 | 27.4 | 0.9909 | Likely Pathogenic | VUS |
| 8 | Missense | 16 | 86510915 | c.346C>T | p.(Pro116Ser) | Forkhead DBD | 0.0000758 | Probably damaging | Deleterious | Pathogenic Moderate | Pathogenic Moder | Uncertain | 0.803 | 27.4 | 0.9909 | Likely Pathogenic | VUS |
| 9 | Missense | 16 | 86510915 | c.346C>T | p.(Pro116Ser) | Forkhead DBD | 0.0000758 | Probably damaging | Deleterious | Pathogenic Moderate | Pathogenic Moder | Uncertain | 0.803 | 27.4 | 0.9909 | Likely Pathogenic | VUS |
| 10 | Missense | 16 | 86512998 | c.1053G>T | p.(Met351Ile) | General AD | 0.0000758 | Benign | Tolerated | Benign Supp | Uncertain | Benign Supp | 0.327 | 23.9 | 0.5891 | Likely Pathogenic | VUS |
| 11 | Missense | 16 | 86513077 | c.1132G>A | p.(Val378Met) | General AD | 0 | Probably damaging | Deleterious | Pathogenic Supp | Pathogenic Moder | Uncertain | 0.574 | 32 | 0.5703 | Likely Pathogenic | VUS |
| 12 | Missense | 16 | 86511182 | c.613G>T | p.(Gly205Cys) | Cell Specific AD | 0.0000121 | Probably damaging | Deleterious | Pathogenic Moderate | Pathogenic Supp | PathogenicSupp | 0.542 | 26 | 0.1346 | Likely Benign | VUS |
| 13 | Missense | 16 | 86510805 | c.236A>G | p.(Gln79Arg) | Forkhead DBD | 0.00000561 | Probably damaging | Deleterious | Pathogenic Supp | Uncertain | Pathogenic Moderate | 0.67 | 24.4 | 0.9892 | Likely Pathogenic | VUS |
|  |  |  |  |  |  |  | 0 |  |  |  |  |  |  |  |  |  |  |

Abbreviations: Chr: chromosome; Cat: category; Supp: supporting; DBD: DNA binding domain; ACMG: American College of Medical Genetics; VUS: variant of unknown significance; LB: likely benign, PS: Pathoenicity score, inor allele frequency using gnomAD genomes version Version: 3.1.2

### Frequency of FOXF1 Variants in Patients with PAH

To estimate the prevalence of *FOXF1* coding variants in patients with PAH, we employed a tiered-search approach, using exome sequencing genomic data from a large cohort, including 2534 patients enrolled in the National Biologic Sample and Data Repository for PAH (PAH Biobank). After filtering for rare variants (gnomADv2.1.1 minor allele frequency <0.01), three *FOXF1* coding variants were identified in five patients with PAH, distinct from the variant identified in the two index cases described above (**Table 2**). These patients did not carry mutations in genes previously associated with PAH including *BMPR2, ACVRL1, ENG, SMAD1, SMAD4, SMAD9, KCNK3, TBX4, EIF2AK4, AQP1, ATP13A3, GDF2* or *SOX17*. Two additional patients with rare *FOXF1* variants and concomitant pathogenic variants in *BMPR2* and *GDF2* were identified but were excluded from further analysis. To determine if the prevalence of *FOXF1* coding variants was greater than expected for the general population, we compared the *FOXF1* variant burden between the PAHBiobank and gnomADv2.1 datasets. Rare, predicted deleterious *FOXF1* protein coding variants were significantly more frequent per individual in the PAH Biobank compared to control cases in gnomAD v2.1 (*χ*^2^ 29.07, *p <* 0.00001). However, no significant difference in synonymous variants was identified (*χ*^2^ = 1.82, p = 0.18).

**Table 2.**
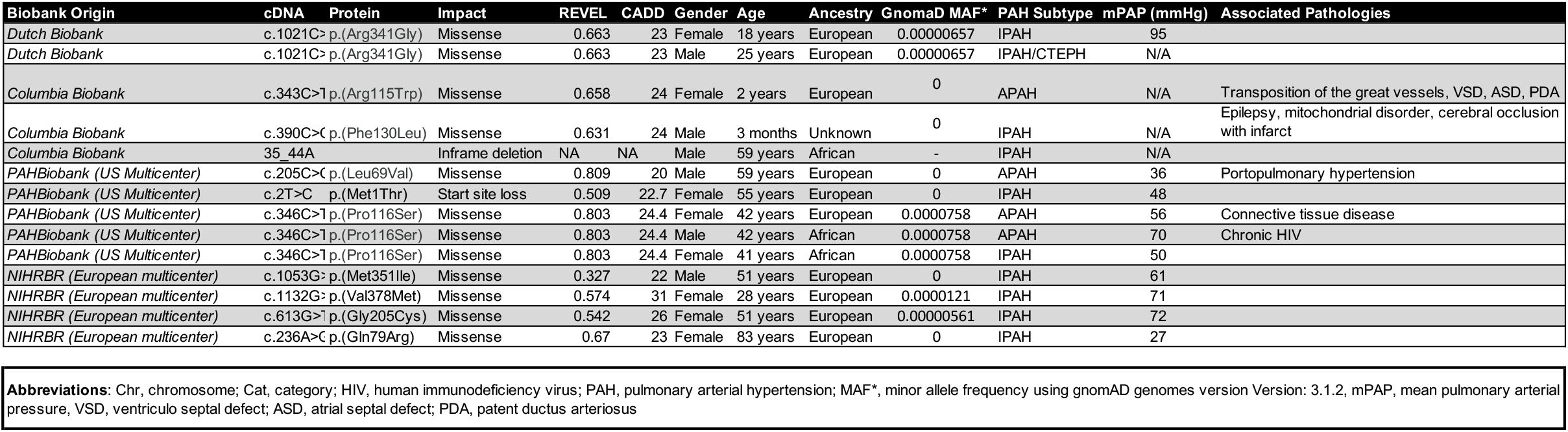
Clinical and demographic information of PAH carrying FOXF1 rare variants.

Like most human transcription factors(37), *FOXF1* exhibits strong genomic regional constrain within the area that encodes the DNA binding domain (chr16:86,510,698-86,511,008, GRCh38/hg38). In essence, the lack of genetic variation within this region implies a strong purifying selection due to essential function or disease pathology(38). Based on homology modeling of *FOXF1*(39, 40), two variants (L69V and P116S, found in 4 different PAH patients) were located within the highly conserved helix-winged-helix DNA-binding core (**Figure 1E-H)**. Importantly, these two variants had an alternative allele frequency of 0.56% in the PAH Biobank cohort, compared to 0.15% in control cases in gnomADv2.1.1, indicating an enrichment in patients with PAH (relative risk 3.73, *p<0*.*0001*).

To validate our findings, we conducted a separate tiered search for rare *FOXF1* variants (CADD scores >19) in two independent genomic datasets from the NIHRBR and CUIMC. Seven additional *FOXF1* variants were found in eight patients, including four novel and three rare variants, as well as an in-frame deletion of a single amino acid. **Figure 1E** displays the distribution of all missense variants within the *FOXF1* gene found in the three datasets, along with their predicted spatial location within the gene (**Figure 1H)**. None of these patients had additional deleterious variants in PAH-risk genes. However, six patients with rare FOXF1 variants in addition to deleterious mutations in *BMPR2, ACVLR1, EIF2AK4* **<u>or</u>** *TBX4*, were identified but excluded from further analysis.

Clinical data, gene and protein changes, allele frequency, estimated protein domain location, and consequence type based on the *in-silico* tools used to determine deleteriousness are listed in **Table 1 and Table 2. Supplemental Table 1** shows ancestry-specific allele frequencies for all variants. In total, 6 of the 11 variants were absent in publicly available allele frequency datasets. **Supplemental Table 2** summarizes the details of all *FOXF1* variants found in patients with concomitant mutations in PAH-risk genes. Phenotypically, patients with *FOXF1* variants were frequently diagnosed with IPAH (73%) and belonged to the Northern European heritage group (73%) (**Table 2**). Four patients had PAH associated with other comorbidities including congenital heart disease, connective tissue disease, chronic HIV infection and portopulmonary hypertension. Of note, none of these *FOXF1* variants have been previously reported in patients with ACDMPV (29). However, other ACDMPV-associated variants involving nucleotides with proximity (1-3 bases), spanning all regions of the *FOXF1* gene, have been reported(29).

Comparative genomic analysis showed that all *FOXF1* missense variants localized to highly conserved genomic sites across eutherian species (**Figure 1I and Supplemental Figure 2)**. We found five variants (p.Leu69Val, p.Gln79Arg, p.Arg115Trp, p.Pro116Ser, p.Phe130Leu) occurring within the forkhead box DNA binding domain (DBD, **Figure 1H**). Variants p.Leu69Val (located in a hinge region between helix 1 and helix 2) and p.Pro116Ser (located within the wing-2 domain) involved residues with high conservation within the DBD among the other 50 members of the FOX family(40) (**Figure 1I**). Although p.Gln79Arg, p.Arg115Trp and p.Phe130Leu occurred in the DBD, these residues were not as highly conserved within the DBD among other FOX members. None of the *FOXF1* variants involved putative DNA binding residues (**Supplemental Figure 3**). **Supplemental Table 3** summarizes complementary *in-silico* analyses to predict the structural or functional effects of all *FOXF1* variants.

**Figure 2.**
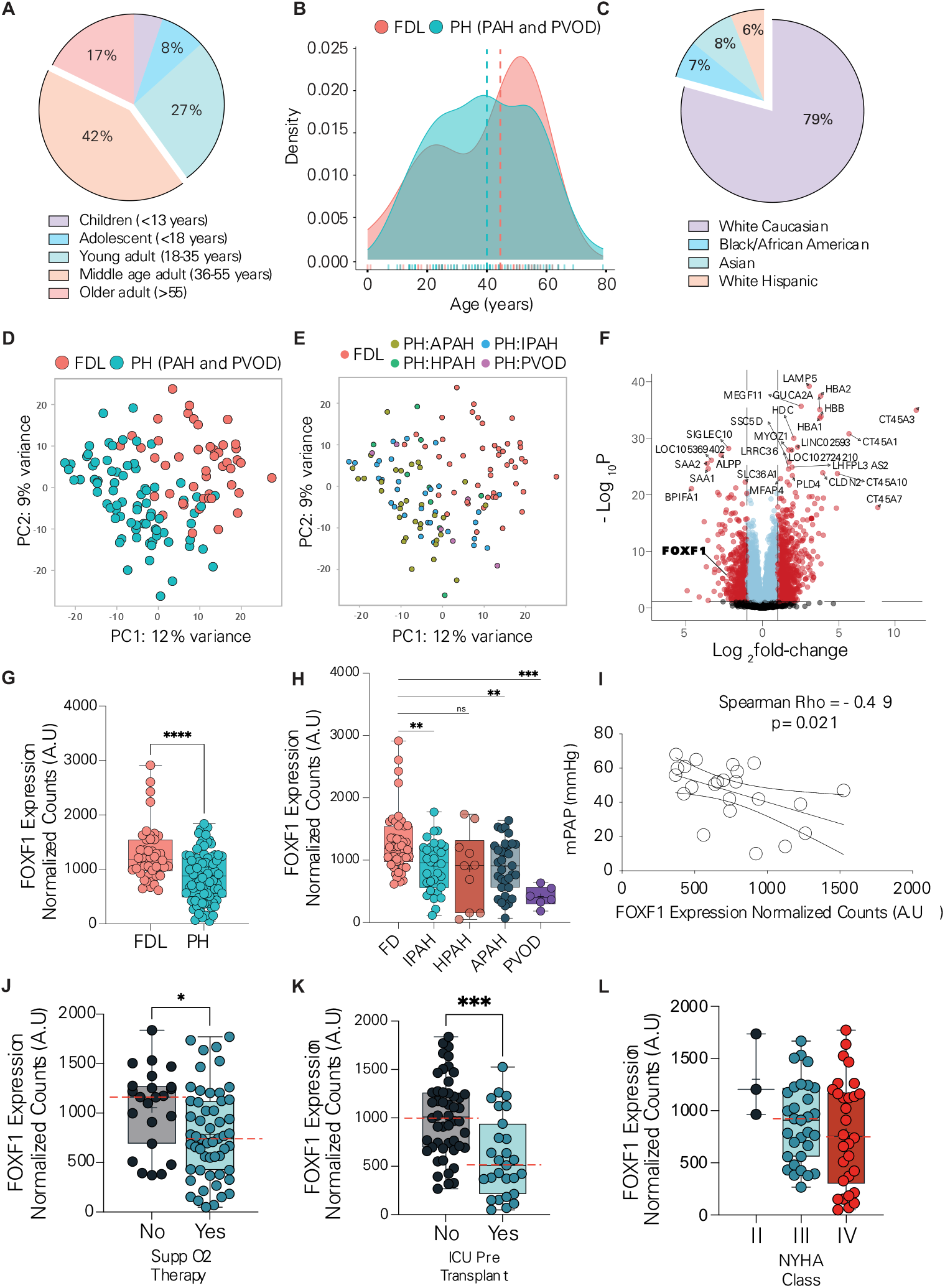
Analysis of bulk RNA-seq data performed on explanted pulmonary arterial hypertension and control lungs. A) Age distribution of subjects in the PHBI biobank, including failed-donor lungs (FDL) and pulmonary arterial hypertension (PAH) patients. B) Density plot showing age differences between FDL and PAH groups. C) Racial distribution of all subjects in the PHBI. D) PCA plot of gene expression profiles illustrating separation between FDL and PAH groups, with no specific heterogeneity within PAH subtypes (E). F) Volcano plot of differentially expressed genes between conditions, highlighting genes with a Log2 fold-change > 0.5 (red). G-H) Box-and-whisker plots of FOXF1 normalized counts between FDL and PAH, including PAH subtypes. The median FOXF1 expression level is significantly lower in PAH samples compared to FDL. I) Correlation of FOXF1 mRNA counts with mean pulmonary artery pressure (mPAP) measured by right heart catheterization at transplantation. Patients exhibiting signs of severe PAH such as the need for supplemental oxygen therapy (J), need for intensive care unit utilization peri-transplant (K) exhibited significantly lower expression of FOXF1. A trend towards lower FOXF1 expression in patients with worse New York Heart Association (NYHA) functional class was observed but did not reach statistical significance. Abbreviations: IPAH = idiopathic PAH, HPAP = hereditary PAH, APAH = PAH associated with other medical co-morbidities, PVOD = pulmonary venooclussive disease. P-value ≤0.05 (*), p-value ≤0.01 (**), p-value ≤0.001 (***), p-value ≤0.0001 (***).

**Figure 3.**
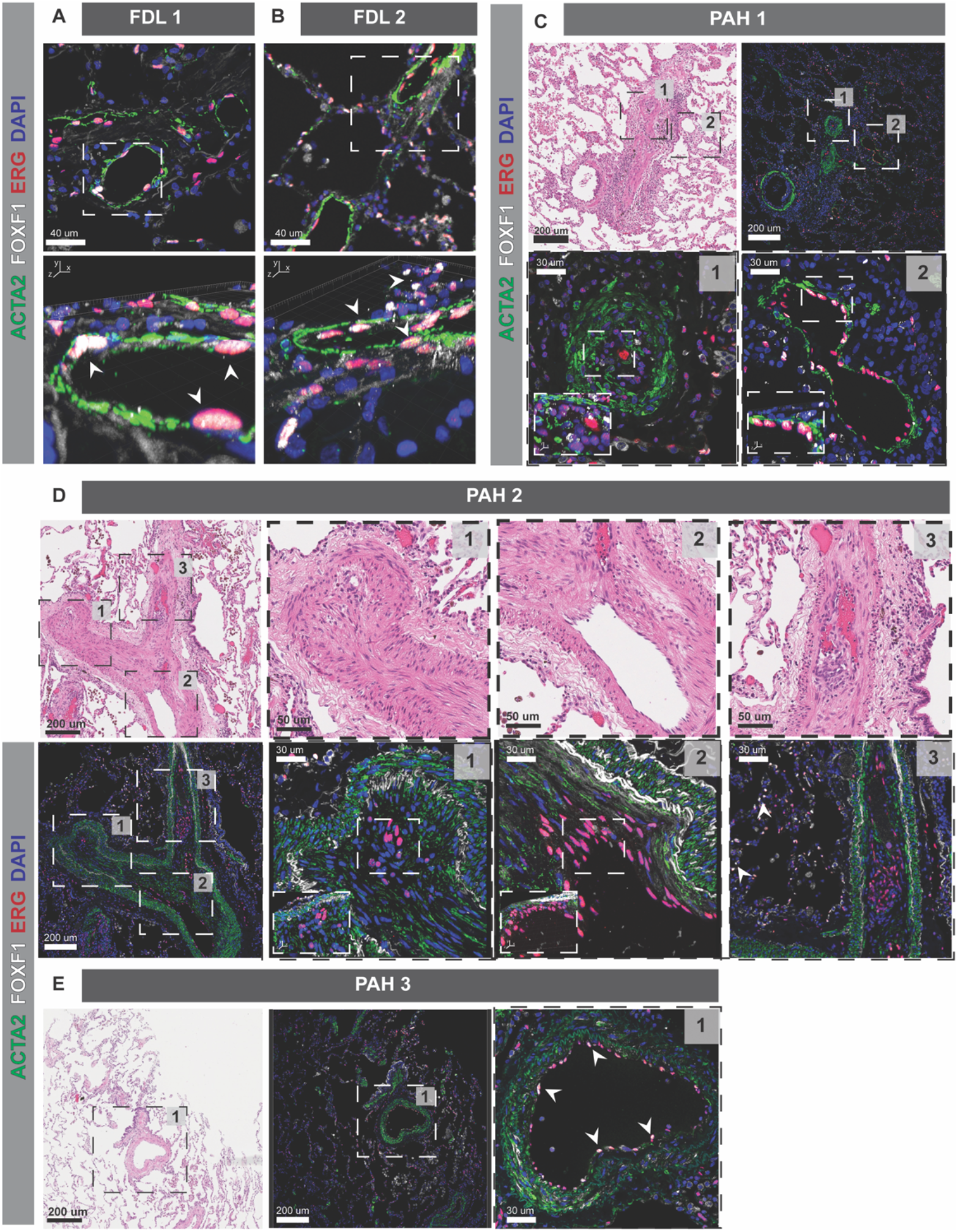
Immunofluorescence analysis of human PAH and control lung tissue samples. High-resolution confocal images demonstrate the co-localized immunofluorescence of FOXF1 (white) with the endothelial marker ERG (red) in failed-donor lung (FDL) controls and pulmonary arterial hypertension (PAH) lung tissues. The smooth muscle alpha (α)-2 actin protein marker (ACTA2) was used to stain vascular smooth muscle cells. DAPI (4’,6-diamidino-2-phenylindole) was used as the fluorescent nuclear counterstain. In FDL tissue, FOXF1 staining is present in arterioles (A) and capillary endothelial cells (B). In contrast, FOXF1 expression appears to be lost in endothelial cells within complex vascular lesions in PAH patients (C-D). FOXF1 expression can still be observed in other regions of the same vascular lesions without overt signs of abnormal cell proliferation (C-2 and D-2). In PAH samples, vessels characterized by muscularization rather than cellular proliferation show endothelial cell FOXF1 staining co-localized with ERG (E). Higher magnifications are marked by small frames. The bottom panels of A-B represent magnified orthogonal views (xyz), where arrows indicate endothelial cell nuclei with co-expression of FOXF1 and ERG. In panels C and D, orthogonal views are shown in small frames. Representative images of vascular lesions are provided with corresponding hematoxylin and eosin staining.

### FOXF1 Expression in Human Lung Tissue and Associated Phenotypic Features in Patients with PAH

In addition to deleterious mutations in the coding regions of *FOXF1*, we postulated that dysregulation of *FOXF1* expression would be associated with the development or progression of PAH. To test this hypothesis, we analyzed human explanted lung tissue samples derived from the PHBI biobank. A total of 135 explants were studied, encompassing 50 failed-donor control lungs (FDL) and 85 samples from PAH patients including idiopathic, hereditary, and associated (secondary) forms of the disease. Additionally, six patients with pulmonary venoocclusive disease (PVOD) were included in the analysis. PHBI samples have been clinically and histologically characterized by others(32, 41, 42). As illustrated in **Figure 2A**, the biobank included tissue derived predominantly from young adults (18-35 years) and middle-aged adults (36-55 years). The median age for PAH patients was 39 years (95% C.I 35-42) compared to 45 years (95% C.I. 35-44) in the FDL group (**Figure 2B**). Female sex was most prevalent in PAH patients (3:1), however males were more prevalent in the FDL group (2:1). As shown in **Figure 2C** most patients self-reported as White. Thirty-four patients had a diagnosis of PAH associated with a comorbidity including congenital heart disease (n=17), connective tissue disease (n=10, scleroderma and systemic lupus erythematosus being the most common) and drug-associated PAH (n=7). Most patients were in functional class III and IV, and approximately 70% of patients were receiving some form of prostacyclin therapy (intravenous or subcutaneous). The median mPAP on diagnostic right heart catheterization was 55 mmHg (95% I.C. 54-63) and the median pulmonary vascular resistance (PVR) was 12.51 (95% C.I. 10-14), suggesting that most patients had severe disease.

Principal component analysis of bulk RNA transcriptome datasets derived from PAH and FDL lung tissue showed a marked shift in the distribution between the groups (**Figure 2D)**. Notably, no specific transcriptional heterogeneity was observed within PAH subtypes (**Figure 2E**). Differential gene expression analysis, including age and sex as co-variables in the model, yielded 10,536 differentially expressed genes (false discovery rate <0.1), out of which 5886 were upregulated and 4650 were downregulated (**Figure 2F**). Age and sex were selected as co-variates due to differences in sex ratios between FDL and PAH groups, as well as the older age in the FDL group. **Supplemental Table 4** includes all differentially expressed genes. Importantly, *FOXF1* expression (normalized mRNA counts) was 31% lower in PAH tissue samples compared to FDL (Log_2_ fold-change -0.525, B-H p-value = 0.001; **Figure 2G**). *FOXF1* mRNA count number was statistically lower in all subtypes of PAH compared to FDL, except for the small number of patients with hereditary PAH (**Figure 2H**). Among patients with known pathogenic variants in PAH-risk genes, 3 patients with *BMPR2* variants and 3 patients harboring *TBX4, ACVRL1* and *EIF2AK* mutations, respectively, showed *FOXF1* expression below the median count in the FDL group (**Supplemental Figure 4A**). We found a significant inverse linear association between mPAP measured at the time of transplant (time of tissue collection) and *FOXF1* normalized counts (Spearman Rho = (-)0.49, p<0.05, **Figure 2I**). PAH patients with high-risk clinical features such as chronic hypoxic respiratory failure (**Figure 2J**) or intensive care unit hospitalization pre-transplantation (**Figure 2K**) had significantly lower lung *FOXF1* mRNA expression. Although not statistically different, a trend towards lower *FOXF1* mRNA counts was also observed in patients with worse New York Heart Association (NYHA) functional class (**Figure 2L**).

**Figure 4.**
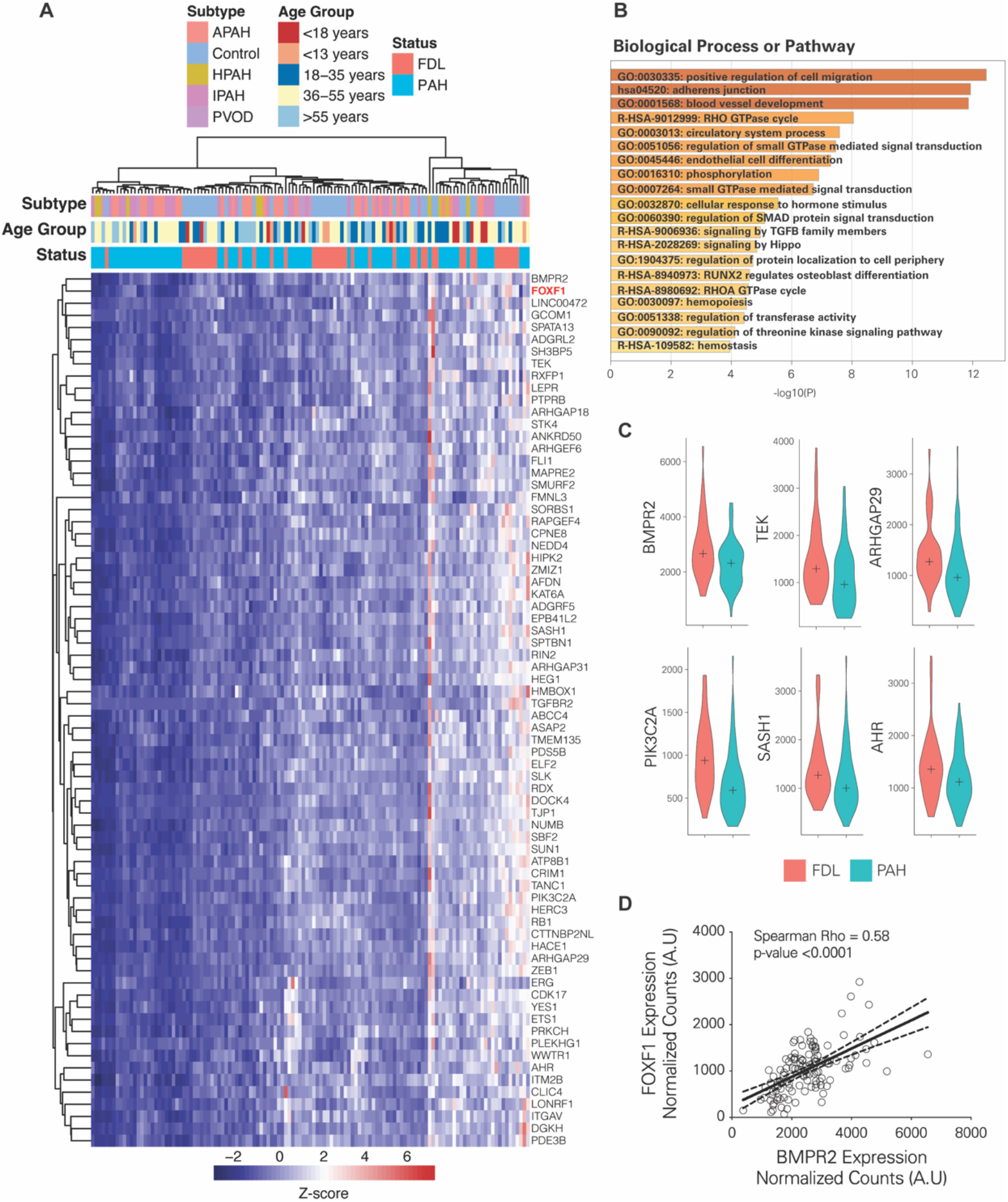
Bioinformatic analysis was conducted using data derived from weighted gene co-expression network analysis on human PAH and control bulk RNA-seq datasets. A) Heatmap showing the expression levels of 72 potential FOXF1 downstream targets across various PAH subtypes and failed-donor control (FDL) samples. Z-scores of gene expression are color-coded from blue (low) to red (high). B) Bar chart of significantly enriched biological processes and pathways. The chart displays -log10(P) values for each pathway, highlighting the top significantly affected biological processes. C) Violin plots illustrating the distribution of normalized mRNA counts for selected FOXF1 targets between FDL and PAH groups. All selected genes are statistically different with a nominal p-value <0.001 and an adjusted p-value (Benjamini-Hochberg False Discovery Rate <0.1). The central cross within each violin represents the median expression level. D) Correlation plot between FOXF1 and BMPR2 normalized mRNA counts. Abbreviations: A.U = arbitrary units, IPAH = idiopathic PAH, HPAP = hereditary PAH, APAH = PAH associated with other medical co-morbidities, PVOD = pulmonary venooclussive disease. Genes: *BMPR2* = Bone Morphogenetic Protein Receptor Type II, *TEK* = TEK receptor tyrosine kinase, *ARHGAP29* = Rho GTPase activating protein 29, *PIK3C2A* = phosphatidylinositol-4-phosphate 3-kinase catalytic subunit type 2 alpha, *SASH1* = SAM and SH3 Domain Containing 1, *AHR* = Aryl Hydrocarbon Receptor.

The long noncoding RNA *FENDRR* shares a promoter with *FOXF1*, and its expression is regulated in *cis* by a *FOXF* lung-specific distant enhancer and int*rans* by *FOXF1*(43). We found a moderate positive correlation between *FOXF1* and *FENDRR* in the PHBI dataset (**Supplemental Figure 4B**), which may indicate transcriptional repression of the *FOXF1/FENDRR* promoter and/or decreased *FOXF1* transcriptional activity in the lungs of patients with PAH.

### Histopathological Features and Distribution of FOXF1 Expression in PAH Lungs

In neonates with ACDMPV, *FOXF1* dysfunction is associated with pathognomonic histological changes such as severe muscularization of small peripheral pulmonary arterioles, thickening of interalveolar septa, reduction of alveolar capillaries and peripheral veins running alongside pulmonary arteries, the latter of which is termed “misalignment of pulmonary veins” (44–48). Many of these features are present in mouse models with *Foxf1* haploinsufficiency(26, 49) or transgenic mice carrying a deleterious DBD knock-in *Foxf1* variant (S52F)(50). In the histological samples available from the PHBI biobank, we did not identify misalignment of pulmonary veins, a key diagnostic feature of typical ACDMPV. In contrast, we observed classical histological features of PAH including extensive muscularization of precapillary pulmonary arteries, medial hypertrophy, endothelial cell proliferation with intimal occlusion, as well as plexiform lesions (**Supplemental Figure 5**).

**Figure 5.**
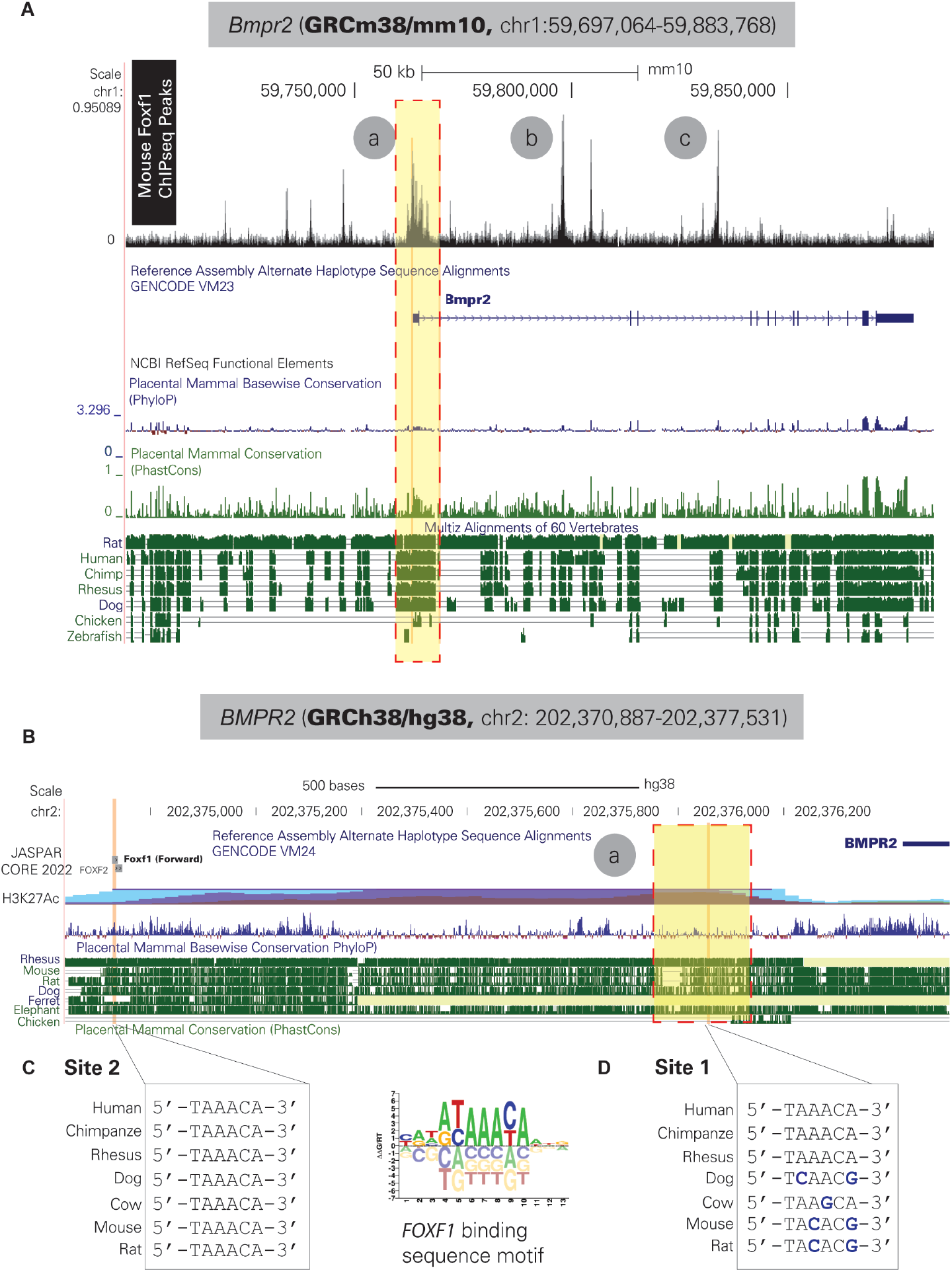
Location of FOXF1 predicted binding sites within mouse and human within the *BMPR2* gene. A) Foxf1-ChIP-seq peaks in relation to the mouse Bmpr2 transcription start site and exon 1 were generated using Cistrome DB(117) from the GSM2062274 data by Dharmadhikari et al. Three peaks (“a”, “b”, “c”) indicate Foxf1 binding sites (ChIP-seq peaks). Peak “a”, located within the Bmpr2 promoter region (shown inside the yellow box), is highly conserved across eutherian species as shown by conservation scores across species (PhyloP and PhastCons tracks)(61). B) Mouse Foxf1 binding peak “a” coordinates, when converted into the human genome assembly, map to a genomic region within the BMPR2 promoter (yellow box), which includes a predicted FOXF1 binding site (Site 1). C) The Site 1 putative binding site is highly conserved among primate species but only partly conserved among other eutherian species. Close to the BMPR2 transcription start site, an alternative FOXF1 binding site was identified, which coincided with a mouse Foxf1-predicted binding site from the JASPAR database (Site 2) and was highly conserved across eutherian species. The gray box shows the genome assembly used to generate the figure, as well as the specific coordinates illustrated. A logo showing the predicted human FOXF1 sequence binding motif was obtained from the Cis-BP database (118).

In FDL samples, we detected nuclear expression of FOXF1 in small arterioles ranging from 30 to 75 micrometers, as well as in capillary EC. In contrast, in PAH lung tissue, nuclear FOXF1 staining was either low or absent within vessels exhibiting disorganized EC proliferation or plexiform lesions (**Figure 3C-D**). While FOXF1 staining was decreased or was absent in regions of vessels undergoing destructive angio-obliteration (**Figure 3D**), we observed FOXF1 staining within distinct regions of the same vessels that had not yet undergone occlusion due to abnormal EC proliferation. Positive staining for ETS-related gene (*ERG*), an endothelial cell marker with nuclear localization similar to FOXF1, indicates that the absence of FOXF1 staining was not a result of nuclear penetration by the antibodies. FOXF1 expression was generally preserved in EC of vessels with increased muscularization but without angio-obliteration (**Figure 3E**). Collectively, these data support the concept that FOXF1 protein expression is decreased in EC within vessels that are remodeled by excessive EC proliferation.

### Weighted Gene Co-expression Network Analysis Identifies FOXF1-Related Genes with Differential Expression in PAH Lungs

To identify genes functionally related to *FOXF1* in PAH human lung tissue, the PHBI bulk RNAseq datasets were used to generate gene co-expression modules and construct scale-free topology networks, using the weighted gene co-expression network analysis (WGCNA) R package (see methods)(51). We identified 50 modules that clustered based on module eigengene, which represents the first principal component of gene expression (51). *FOXF1* clustered with 2902 genes, with various degrees of correlation (**Supplemental Figure 4C**). To define potential downstream genes directly affected by decreased *FOXF1* expression, we crossmatched the 2902-gene module with a list of predicted *FOXF1* targets recently identified through multiomic (ATAC and RNAseq) single cell sequencing of normal and ACDMPV human lung tissue(52). This process led to the identification of 72 potential *FOXF1* downstream targets (**Figure 4A and Supplemental Table 5**). In agreement with previous work, 87% of these genes (68/72) are also differentially expressed in patients with ACDMPV carrying *FOXF1* pathogenic mutations compared to normal lung tissue(53). Based on conserved expression markers, we predicted these genes to be predominantly expressed in CAP1 ECs (general capillary ECs), CAP2 ECs (aerocytes expressing *CAR4*), systemic vasculature ECs (SVEC, also known as peribronchial ECs), pericytes, as well as mesenchymal (AF1, expressing *TCF21*) and alveolar (AF2, expressing *MFAP5*) fibroblasts (**Supplemental Table 5**)(52, 54, 55).

Functional analysis using *Metascape*(56) revealed distinct biological processes and pathways associated with the predicted targets, with the top 20 displayed in **Figure 4B**. Remarkably, we observed a significant enrichment for pathways linked to the positive regulation of cell migration (GO:0030335), blood vessel development (GO:0001568), EC differentiation (GO:0045446), and signaling by TGFB family members (R-HSA-9006936). *Metascape* analysis indicated that 18/20 Top Cell type signatures were enriched for EC datasets from various organs including kidney, heart, lung, and liver (**Supplemental Figure 4D**). Altogether, these data demonstrate that the *FOXF1*-related transcriptional signature includes targets with the ontological associations expected for a transcription factor that is essential to blood vessel development.

We next utilized *DisGeNET* (57) to compare the *FOXF1*-related transcriptional signature with various data repositories including GWAS catalogues, scientific literature, and animal models of different human diseases. As shown in **Supplemental Figure 4E**, the strongest association was found with “pulmonary arterial hypertension” (C3203102). Other interesting signatures included “scleroderma” (C0011644), “hemangiosarcoma” (C0018923) and different types of sarcoma-related gene ontologies. Only 38% of *FOXF1* predicted downstream genes (28/72) were included in the list of differentially expressed genes between FDL and PAH lung tissue samples (**Supplemental Table 6**). Using this 28-gene list, *Metascape* analysis identified pathways and biological processes enriched for positive regulation of cell migration (GO:0030335), blood vessel development (GO:0001568) and endothelial cell differentiation (GO:0045446). In addition, we found several genes involved in protein ubiquitination (*HACE1, HERC3, NEDD4*) and several tumor suppressor genes (*CLIC4, HIPK2, NUMB, RB1*). We selected six specific target genes based on predicted EC expression in human lung tissue from single cell sequencing atlases(54, 58), their potential relevance to PAH and their expression difference between PAH and FDLs (**Figure 4C)**. Most importantly, we identified *BMPR2* as a putative *FOXF1* binding target in human lungs. *BMPR2* mRNA counts were statistically lower in patients with PAH (**Figure 4C**), and significantly correlated with *FOXF1* RNA levels (**Figure 4D)**.

To orthogonally validate our results from human data, we used publicly available mouse E18.5 lung FOXF1 ChIP-seq datasets(59) and assessed the presence and distribution of predicted *FOXF1*-binding sites within the *Bmpr2* gene region. A modified UCSC Genome Browser(60) screenshot overlaying murine FOXF1 ChIP-seq peaks with the GENCODE transcript set (Release M23/GRCm38.p6), and comparative genomics (placental conservation) tracks is shown **Figure 5**. Three sites directly bound by FOXF1 protein were identified (**Figure 5A**); one of these peaks (chr1:59,763,118-59,763,467) was located within 1 kb of the *Bmpr2* transcription start site (TSS), suggesting a role for FOXF1 in binding the *Bmpr2* promoter. We converted the mouse genomic coordinates to the human genome assembly GRCh38/hg38 (chr2:202,375,753-202,376,331) and identified two putative biding sites located within 1 kb of the *BMPR2* TSS (**Figure 5B**). Putative site 1 (chr2:202,375,956-202,375,961) is located within the predicted *BMPR2* promoter sequence. Putative site 2 (chr2:202,374,830-202,374,836) was located within 1.5 kb of the *BMPR2* TSS but outside of the predicted *BMPR2* promoter region (**Figure 5C-D**). Site 2 was evolutionarily conserved among eutherian species and aligned with a mouse FOXF-binding site according to JASPAR(61) and TRANSFAC(62)) data. However, the site 1 sequence was only identical within primate species (data not shown) but showed on average a two-nucleotide difference with other eutherian species. In addition to the location within the BMPR2 promoter (Site 1) and the proximity to the TSS (Site 2), both predicted FOXF1 binding sites are located within a genomic region enriched for the H3K27Ac histone mark in human umbilical vein endothelial cells (HUVEC), normal human lung fibroblasts (NHLF), and human embryonic stem cell lines (H1-hESC), suggesting that these sites are located in areas of enhanced transcription in these cell types (**Figure 5B;** “H3K27Ac track”). Collectively, these data suggest that *BMPR2* represents a potential FOXF1 downstream target gene, based on co-expression analysis, differential expression between PAH and FDLs and the presence of putative FOXF1 binding sites (via ChIP-seq and predicted FOXF1 binding sequences) within regulatory regions directly upstream of the *BMPR2* gene.

## Discussion

The discovery of the genetic basis for PAH revolutionized our understanding of the disease and led to a new generation of BMPR2-centric pharmacotherapies (63, 64). While most studies in PAH genetics have focused on identifying protein-truncating variants (i.e., protein-truncating variants)(19), the recent discovery of mutations in *KDR*(31) provided the framework for a tiered, hypothesis-driven approach to identify ultra-rare missense variants associated with the disease. Using a similar approach, we identified novel and rare *FOXF1* coding variants in different multicenter cohorts of patients with PAH. In parallel, we demonstrate that *FOXF1* expression is dysregulated in the lungs of patients with PAH, and this dysregulation is associated with disease severity. *FOXF1* mRNA levels strongly correlated with *FENDRR* expression, a long non-coding RNA that is divergently transcribed from a shared promoter and co-expressed with *FOXF1*, suggesting that transcriptional repression may contribute to *FOXF1* downregulation in the lungs of PAH patients. Using histological samples, we showed that *FOXF1* expression is heterogeneous, with the lowest or absent expression in phenotypically abnormal EC within complex vascular lesions. In the samples available to us, we did not find evidence to support that low *FOXF1* expression was associated with histological features of ACDMPV. Lastly, using a hybrid, partly-supervised bioinformatic approach, we identified different *FOXF1* downstream targets with potential involvement in the pathogenesis of PAH, including *BMPR2*. Collectively, by leveraging large, independent human genomic and transcriptomic datasets, we provide evidence associating decreased expression or predicted dysfunction of *FOXF1* with a phenotype of PAH.

The Forkhead box (FOX) proteins are a group of evolutionarily conserved transcription factors with remarkable functional diversity in eukaryotic development(27, 65, 66). FOXF1, specifically, plays a crucial role in the normal development of the lungs, liver, and gut(26). In the lung, FOXF1 stimulates angiogenesis and vascular repair after injury by promoting EC proliferation while maintaining vessel integrity(27, 67), via regulation of *VEGF, PDGF, NOTCH2* and *TIE2*(26, 28). Despite this understanding, the precise mechanisms underlying *FOXF1* deficiency in human diseases remain incompletely understood. Research by Wang *et al*. has shown that decreased activin receptor-like kinase 1 (*ACVRL1*) signaling partly explains the ACDMPV phenotype in mice with a pathological *FOXF1* variant (p.Ser52Phe)(68). However, additional mechanisms are likely involved, as mutations in *ACVRL1*(69) (or its ligand, *BMP9*(70, 71)) more commonly manifested in adults with PAH associated with hereditary hemorrhagic telangiectasia, rather than ACDMPV. Recently, *FOXF1* has been implicated in transgenic mice with EC-specific deletion of *Bmpr2*(72), however, its role in human adult PAH remains largely unknown.

Heterozygous missense variants or copy number variation deletions in the *FOXF1* gene on chromosome 16q24.1 are most frequently associated (in 80%–90% of patients) with a phenotype of ACDMPV, a rare, congenital lung disorder characterized by severe respiratory failure and fulminant PAH shortly after birth(29, 73), though several cases with a late presentation of ACDMPV have been reported(47, 74–77). Histological studies from late-survivors with ACDMPV suggest that the variable phenotype can be partially explained by a heterogeneous, non-uniform distribution of parenchymal and vascular changes(47). Whereas it has been postulated that histological heterogeneity may result from somatic mosaicism(48), it is unclear whether the severity of parenchymal or vascular changes is linearly dependent on *FOXF1* expression or its transcriptional activity. Here we propose that the variants identified in PAH patients result in a partly dysfunctional, but not inactive, protein compared to patients with variants resulting in a more severe phenotype, such as ACDMPV. Whereas most variants located in the DNA binding domain ([DBD], the most conserved region of FOXF1) are pathogenic(30), we postulate that the DBD variants found in patients with PAH are hypomorphic because they do not involve residues with high conservation among other FOX members (i.e. FOXC1, FOXA1, etc), do not involve predicted DNA binding residues or they do not severely alter protein conformation. Variants located in disorganized regions outside the DNA binding domain (DBD), may affect protein-protein interactions, thereby impacting the transcriptional function of FOXF1. Variants p.Arg341Gly, p.Met351Ile, and p.Val378Met are predicted to be located in disordered regions outside the DBD. Although the specific functions of these FOXF1 domains are only partly understood(78), many pathogenic variants in these disordered regions have been observed in ACDMPV, indicating their importance in FOXF1 biology(29).

Another explanation for the later age of onset of patients harboring *FOXF1* variants may be the presence of regulatory elements or alternative modifiers. Strikingly, cases of adults carrying pathological *FOXF1* variants have been reported in unrelated families with ACDMPV (29, 48, 79–82). Some of these adult carriers appear to be asymptomatic; however, a deep clinical, radiological or physiological characterization is unavailable. A subset of these “asymptomatic” carriers may have additional compensatory variants in regulatory regions that ameliorate the phenotype, perhaps allowing the subjects to survive until adulthood. Bölükbaşı *et al*. reported a hyperfunctional variant in a *FOXF1* distant enhancer capable of compensating for a *FOXF1* frameshifting variant in a mutigenerational family with ACDMPV(79). We did not observe the same variant (rs560517434-A) within the enhancer region (chr16:86219847-86220047; hg38) in the few patients with whole-genome data available. However, due to the lack of intronic information in the PAHBiobank and CUIMC biobank, we were unable to investigate potential compensatory variants in the rest of the samples.

The observation that our index cases had a parent carrying the same *FOXF1* variant without showing signs of the disease is consistent with the complex regulation and inheritance patterns of *FOXF1*(39). Indeed, similar findings have been shown in patients with mutations in *BMPR2*, where non-penetrance is best characterized, as well as other developmental genes linked to PAH, such as *TBX4*(21) and *KDR*(83). Segregation analyses have revealed that not all carriers of *TBX4* or *KDR* variants develop PAH. Patients with pathogenic *TBX4* variants develop PAH at various ages, exhibiting a broad range of clinical presentations(84), including severe respiratory failure and pulmonary hypertension similar to patients with acinar dysplasia(85). Notably, some individuals with *TBX4* mutations experience severe pulmonary hypertension during neonatal stages, which resolves early but reappears in infancy or early childhood(86). This pattern suggests a potential influence of environmental factors or adaptations to growth and physiological changes in the development of PAH. A biphasic presentation may also be possible for individuals with *FOXF1* variants.

The differential gene expression of *FOXF1* between controls and PAH patients, along with the heterogeneity of expression among different PAH patients, and the correlation between *FOXF1* and *FENDRR* transcript levels suggest the presence of modifiers of *FOXF1* transcription. While the exact transcriptional regulators of *FOXF1* remain unknown, putative effectors such as sonic hedgehog (SHH) and downstream proteins such as GLI Family Zinc Finger 1 (GLI1) are known to be upstream of *FOXF1* during development(39). Tamura *et al*. found that *FOXF1* expression is induced upon DNA damage in a p53-dependent manner and reported a p53 response element within the *FOXF1* gene(87). Isobe *et al*. demonstrated that EC-specific *Bmpr2* knock-out mice exhibited lower levels of *Foxf1*, after exposure to hypoxia(72). However, it is not yet determined whether *FOXF1* is directly downstream of *BMPR2* or if its downregulation is a consequence of *BMPR2* deficiency.

Using transcriptomic and publicly available epigenomic datasets, we identified genes functionally related to *FOXF1*, highlighting *BMPR2*. Mouse FOXF1 ChIP-seq datasets revealed a putative binding site near the *Bmpr2* transcription start site, and in proximal intergenic regions. Genomic coordinates conversion identified similar potential target sites within the human *BMPR2* promoter, highly conserved among primate species. Although a significant association was found between *BMPR2* and *FOXF1* expression, this association cannot determine if *FOXF1* is decreased due to *BMPR2* downregulation or *viceversa*. Interestingly, in contrast to data from *Bmpr2* knock-out mice, not all patients harboring a pathogenic *BMPR2* variant exhibited lower *FOXF1* expression. Indeed, potential bidirectionality is of particular interest, given that *FOXF1* gene replacement (forced expression using viral(72) or non-viral vectors(88)) could potentially be used to restore *BMPR2* expression without the potential negative side effects associated with other transcriptional activators of BMPR2. Conversely, understanding whether transcriptional activators of *BMPR2* (i.e. recombinant BMP9(89)) or activin A ligand traps could impact the phenotype in patients with ACDMPV could have significant implications for this group of patients without a cure.

It has been proposed that EC in PAH exhibit a cancer-like phenotype(90). Our analysis of histological samples, while limited by sample number, suggests that lower or absent FOXF1 is mainly found in EC of vessels already affected or evolving into a disease state. *FOXF1* prevents DNA re-replication to maintain genomic stability in a p53-p21^WAF1^-dependent manner, by activating DNA damage response protein kinases ATM (ataxia telangiectasia mutated) and ATR (ataxia telangiectasia and Rad3 related)(91). In cancer cell lines, silencing *FOXF1* promotes DNA re-replication by disrupting replication stringency, potentially leading to cellular transformation and tumorigenesis(91, 92). Studies have also shown that EC-specific *Atm* knock-out mice exhibit persistent elevated pulmonary pressures after chronic hypoxia exposure, and silencing *FOXF1* in pulmonary artery EC can induce DNA damage(72). Collectively, these observations raise the possibility that *FOXF1*, functioning as a tumor suppressor gene, may play a role in the development of PAH. Future studies on the effect of *FOXF1* haploinsufficiency on the DNA damage response of EC and its role in acquiring a malignant-like phenotype are warranted but fall outside the scope of this study.

## Limitations and future directions

As for many newly discovered genes involved in hereditary diseases, we have not yet investigated if haploinsufficiency or transcriptional repression of FOXF1 is sufficient to cause a PAH phenotype, alone or in association with BMPR2 dysfunction. The use of gnomAD as a control cohort can be considered as a limitation of the study, however many other studies using a similar approach have validated pathogenic targets in PAH(19). On the horizon is the use of whole genome sequencing, which could enable the identification of variants in non-coding regions working as modifiers of FOXF1 transcription in patients with PAH.

## Conclusions

Our data support the hypothesis that reduced *FOXF1* gene expression or protein dysfunction due to variants in key regions of the *FOXF1* gene contributes to the pathogenesis of PAH. Similar to other PAH-risk genes, incomplete penetrance suggests the involvement of modifiers. Future studies are needed to understand the mechanisms underlying FOXF1-dependent endothelial cell function/dysfunction in PAH.

## Acknowledgements and author’s contributions to the study

Conception and design of this study and creation, revision, and final approval of this manuscript: J.G-A, W.C.N, L.M, V.K.K. Analysis and interpretation: J.G-A, W.C.N, L.M. Data acquisition: All authors. Drafting the manuscript for important intellectual content: All authors. Special thanks to Dr. Anne Katrine Johansen, Dr. Norbert F. Voelkel, Dr. Guolun Wang for their critical comments on the project. We thank all of the patients who contributed samples and data to the National Biological Sample and Data Repository for PAH as well the Investigators and Coordinators/Nurses as the Enrolling Centers. The authors would like to thank the Bioimaging and Analysis Facility (especially director Matt Kofron), Cincinnati Children’s Research Foundation for technical support, as well as the CCHMC Pathology Research Core for tissue processing and embedding.

## Methods

### Genomic PAH cohorts

Three independent cohorts were used for our study: 1) United States National Biologic Sample and Data Repository for PAH (PAH Biobank) housed at Cincinnati Children’s Hospital Medical Center (CCHMC) (23, 31, 93, 94); 2) UK National Institute for Health Research BioResource (NIHRBR) Rare Diseases (including 847 patients with PAH) (20, 94); and 3) Columbia University Irving Medical Center (CUIMC)(21, 95, 96). All patient samples were obtained with written informed consent and approval from local institutional review boards. Sample processing and genomic sequencing were performed as previously reported (21, 31, 93, 94). For PAH Biobank, exome sequencing was performed in collaboration with the Regeneron Genetics Center (RGC) or at CCHMC DNA Sequencing and Genotyping Core. Genomic DNA processed at the RGC was prepared with a customized reagent kit from Kapa Biosystems and captured using the SeqCap VCRome 2 exome capture reagent or xGen lockdown probes. Patient DNA samples sequenced at CCHMC were prepared with the Clontech Advantage II kit and enriched using the SeqCap EZ exome V2 capture reagent. PAH Biobank samples were sequenced using an Illumina HiSeq 2500 platform, generating 76-bp paired-end reads. Read-depth coverage was ≥15X in ≥95% of targeted regions for all exome sequencing samples. Burrows-Wheeler Aligner (BWA-MEM) was used to map and align paired-end reads to the human reference genome (GRCh38)(97). The *MarkDuplicates* from the Picard package was used to identify and flag polymerase chain analysis duplicates, and the *HaplotypeCaller* from GATK (genome analysis toolkit) was used to call genetic variants(98).

For the UK cohort, DNA extracted from venous blood underwent whole-genome sequencing using the Illumina TruSeq DNA PCR-Free Sample Preparation kit (Illumina Inc., San Diego, CA, USA) and Illumina HiSeq 2000 or HiSeq X sequencer, generating 100–150 bp reads with a minimum coverage of 15× for ∼95% of the genome (mean coverage of 35×). Whole-exome sequencing was conducted for individual II-1 using genomic DNA extracted from peripheral blood. Paired-end sequence reads were generated on an Illumina HiSeq 2000. Specific details on patient selection and composition, as well as generation of analysis-ready datasets for the NIHRBR cohort can be found in Gräf *et al*(20).

For Columbia University Irving Medical Center, the whole exome sequencing libraries were prepared with the Agilent SureSelect Paired-End Version 2.0 Human Exome Kit (Agilent, Santa Clara, CA). Sequencing of post-enrichment shotgun libraries was performed on an Illumina Genome Analyzer II following manufacturer’s protocol for 50 bp paired-end reads. Specific details on patient selection and composition, as well as generation of analysis-ready datasets for the CUIMC cohort can be found in Ma *et al*(96).

### Identification of deleterious variants

Population allele frequency was derived from public databases including Exome Aggregation Consortium (ExAC)(99) and Genome Aggregation Database (gnomADv2.1.1)(34). Rare variants were defined by an AF < 0.01% in both ExAC and gnomADv2.1.1 exome sequencing samples (European and non-European) datasets.

First, we selected all non-synonymous *FOXF1* variants using a Combined Annotation Dependent Depletion (CADD) Phred-scale score of >20 <u>or</u> a Rare Exome Variant Ensemble Learner (REVEL) score of >0.5. To aid in variant selection, we determined the median (25th-75th percentile) CADD score for 29 different *FOXF1* deleterious missense variants reported in ClinVar(100), including 18 variants previously published by Sen and Stankiewicz(80). The mean score for these variants was 26.8 (24-31), which was similar to the variants selected in our PAH population with a median (25th-75th percentile) of 25 (22-27).

To estimate deleteriousness, we used various composite scores including metaSVM(101), metaRNN(102) and M-CAP (103) and AlphaMissense(104). These various prediction models integrate information from different high-level annotation scores including functional prediction scores, conservation scores and allele frequency information from the 1000 Genomes Project, ExAC, gnomAD-exome, as well as gnomAD-genomes. Specifically, AlphaMissense, utilizes a deep learning model that builds upon the protein structure prediction tool AlphaFold2, and it is trained on population frequency data.

None of the variants selected were detected in the rare disease controls in NIHRBR or gnomADv2.1.1. The Varsome online platform (www.varsome.com) was used to summarize all data(105). Based on American College of Medical Genetics and Genomics guidelines(106, 107), variants were identified were classified as pathogenic, likely pathogenic, variant of uncertain significance (VUS), likely benign and benign.

Pathogenic variants in established PAH genes have been defined and validated previously(19, 23, 93, 95, 96, 108–110). Samples were screened for variants in known risk genes for PAH including *ACVRL1, BMPR2, CAV1, EIF2AK4, ENG, KCNK3, SMAD4, SMAD9, and TBX4*. The data from the US PAH Biobank and the CUIMC are available via an application-based process.

### FOXF1 variant enrichment analysis

To determine whether individuals with PAH were enriched for protein-coding *FOXF1* variants compared to the general population, we conducted gene enrichment analyses. Due to data availability, these analyses were restricted to the PAH Biobank. First, we identified all rare (minor allele frequency [MAF] < 0.01) protein-coding variants within the PAH Biobank cohort and compared them to those in gnomAD v2.1.1 (n = 141,389). Since gnomAD v2.1.1 aggregates data from multiple studies employing different sequencing platforms (including both exome and genome sequencing), there is variability in the number of participants with called variants. To account for this variability, we restricted the analysis to variants that were called in at least 80% of the gnomAD samples. Next, we performed chi-square goodness-of-fit tests to compare the number of individuals with an alternative allele versus those without the alternative allele for protein-coding variants in *FOXF1*. To control for potential biases between PAH participants and the gnomAD controls, we also conducted similar analyses focusing on rare synonymous variants.

### Protein sequence and structure analysis

The 3D protein model of the reference FOXF1 was retrieved from the AlphaFold2 database(111) using the UniProt ID Q12946 (AlphaFold identifier AF-Q12946-F1). All 3D renderings were generated using PyMol (The PyMOL Molecular Graphics System, Version 1.2r3pre, Schrödinger, LLC). Secondary structure and amino acid conservation analyses were conducted utilizing the POLYVIEW-2D server(112). Putative DNA binding sites were identified using the Conserved Domains Database (CDD).

### Bulk RNA sequencing, outlier filtering and differential expression analysis and data availability

Information on patient enrollment, as well as the tissue-processing protocol for Pulmonary Hypertension Breakthrough Initiative (PHBI) have been previously described(32), and can be found at *http://phbi.org*. For bulk RNAseq, paired-end 75 base-pair total RNA libraries were generated for all available PHBI lung samples using an Illumina sequencer. Samples were sequenced in two batches, with a sequencing depth of 20-25 million reads per sample in batch number one and 15-20 million reads per sample in batch number two. Reads were pseudo-aligned and quantified using an index transcriptome version of the GRCh38.p14 human genome (RefSeq GCF_000001405.40) using *Kallisto* with standard settings(113). Transcript-level abundance estimates were imported and summarized as a counts matrix using *tximport*(114). Raw counts were normalized using DESeq2 “median of ratios method”, which considers both sequencing depth and RNA composition (114–116), for differential gene expression analysis. Potential outliers and batch effects of different covariates (i.e. sequencing batch, sex and age) were assessed by hierarchical clustering and principal component analysis. Two patients with WHO Group IV pulmonary hypertension (chronic thromboembolic pulmonary hypertension) were excluded for the analysis *a priori*, given that the pathobiology is significantly different from PAH (Group 1 PH). Five samples (2 from failed-donor lung and 3 from PAH groups) were identified as outliers by hierarchical clustering and principal component analysis (**Supplemental Figure 1A-B**). These samples included a 55-year-old failed-donor lung and a 14-year-old male patient with IPAH that have been identified as outliers by others (41). These samples had a more than 2 standard deviations lower or higher uniquely aligned reads which could explain their abnormal clustering with other samples. Sequencing data were adjusted and corrected for potential batch effects using the *sva* package (v 3.50.0). Differentially expressed genes were selected based on a Benjamini and Hochberg false discovery rate of <0.1 using DEseq2. This threshold was selected to increase the probability of finding FOXF1 downstream binding targets. The heatmap was generated using the *pheatmap* R package and the volcano plot xswas generated using the *EnhancedVolcano* R package (https://github.com/kevinblighe/EnhancedVolcano).

The PHBI sequencing data discussed in this publication have been deposited in NCBI’s Gene Expression Omnibus and are accessible through GEO Series accession number GSE254617 (https://www.ncbi.nlm.nih.gov/geo/query/acc.cgi?acc=GSE254617).

### Weighted Gene Co-Expression Network Analysis (WGCNA) and Identification of FOXF1 transcriptional network

The batch-corrected, normalized gene expression matrix was again evaluated for potential outlier samples by hierarchal clustering using different features of the WGCNA package(51). For downstream analysis, we used the WGCNA v. 1.72-1 R package to identify modules of *FOXF1* co-expressed genes in PHBI lung RNA-seq samples. We used the *pickSoftThreshold* to determine the power value optimized for the highest scale-free topology index (R2) and lowest mean connectivity. A soft-thresholding power of 9 was selected based on R2 > 0.85 and mean connectivity (k = 200). Data was then transformed into a Topological Overlap Matrix (TOM) and a corresponding dissimilarity matrix was calculated. *blockwiseModules* was used for blockwise module analysis, with the following parameters: TOMType = “signed”, power = 9, minModuleSize = 30, mergeCutHeight = 0.25. Module eigengenes representing the first principal component in the dataset were calculated using the *moduleEigengenes* function of WGCNA. To identify a *FOXF1* co-expression network, we identified which module was assigned to *FOXF1*. The list of genes associated/co-expressed with *FOXF1* was then cross-referenced with genes known to have a *FOXF1* binding sites based on ChIP-seq (59), as well as single nuclei ATAC-seq datasets from controls and from patients with ACDMPV (53).

### Histology and Immunofluorescence

Five micrometre (5 µm) thick paraffin embedded sections of 10% formalin fixed human lung tissue were melted at 60°C for two hours, following rehydration through xylene and alcohol, and finally in PBS. Tissue sections were stained with H&E according to standard protocols. For immunofluorescence, antigen retrieval was performed in 0.1 M citrate buffer (pH 6.0) by microwaving. Slides were blocked for 2 hours at room temperature using 4% normal donkey serum (Jackson Immuno Research Laboratories) in PBS containing 0.2% Triton X-100, and then incubated with primary antibodies diluted in blocking buffer for approximately 16 hours at 4°C. Primary antibodies included ACTA2 (1:2000; A5228, Sigma-Aldrich), FOXF1 (1:50, AF4798, R&D Systems), ERG (1:100,AB92513, Abcam). Appropriate secondary antibodies conjugated to Alexa Fluor 488 (Thermo Fisher Scientific, A21202), Alexa Fluor 568, (Thermo Fisher Scientific, A11057) or Alexa Fluor 633/647 (Thermo Fisher Scientific, A31573) were used at a dilution of 1:200 in blocking buffer for 1 hour at room temperature. Nuclei were counterstained with DAPI (1 μg/ml), (D21490, Thermo Fisher Scientific). Sections were mounted using ProLong Gold antifade reagent, (P36930, Thermo Fisher Scientific) mounting medium with coverslip.

### Additional Statistical Analysis

Since the mRNA data were based on counts, non-parametric analyses were performed. Continuous data were visualized using box-and-whisker plots, violin plots, or scatter plots. Differences between groups using isolated *FOXF1* mRNA counts were determined with a Mann-Whitney test. For comparisons involving more than two groups, pairwise comparisons were performed using the Kruskal-Wallis test. To evaluate the relationship between two continuous variables, Spearman correlation was used. Apart from the differential gene expression analysis mentioned above, all single-gene comparisons and correlations with clinical variables were two-sided, with statistical significance set at p < 0.05. The analyses were conducted using IBM SPSS version 28.0 (Armonk, NY), GraphPad Prism version 10 (Boston, MA), and R Studio (PBC, Boston, MA), utilizing various R-bioinformatic packages as detailed in each section.

## Supplemental Figures

**Supp. Figure 1:**
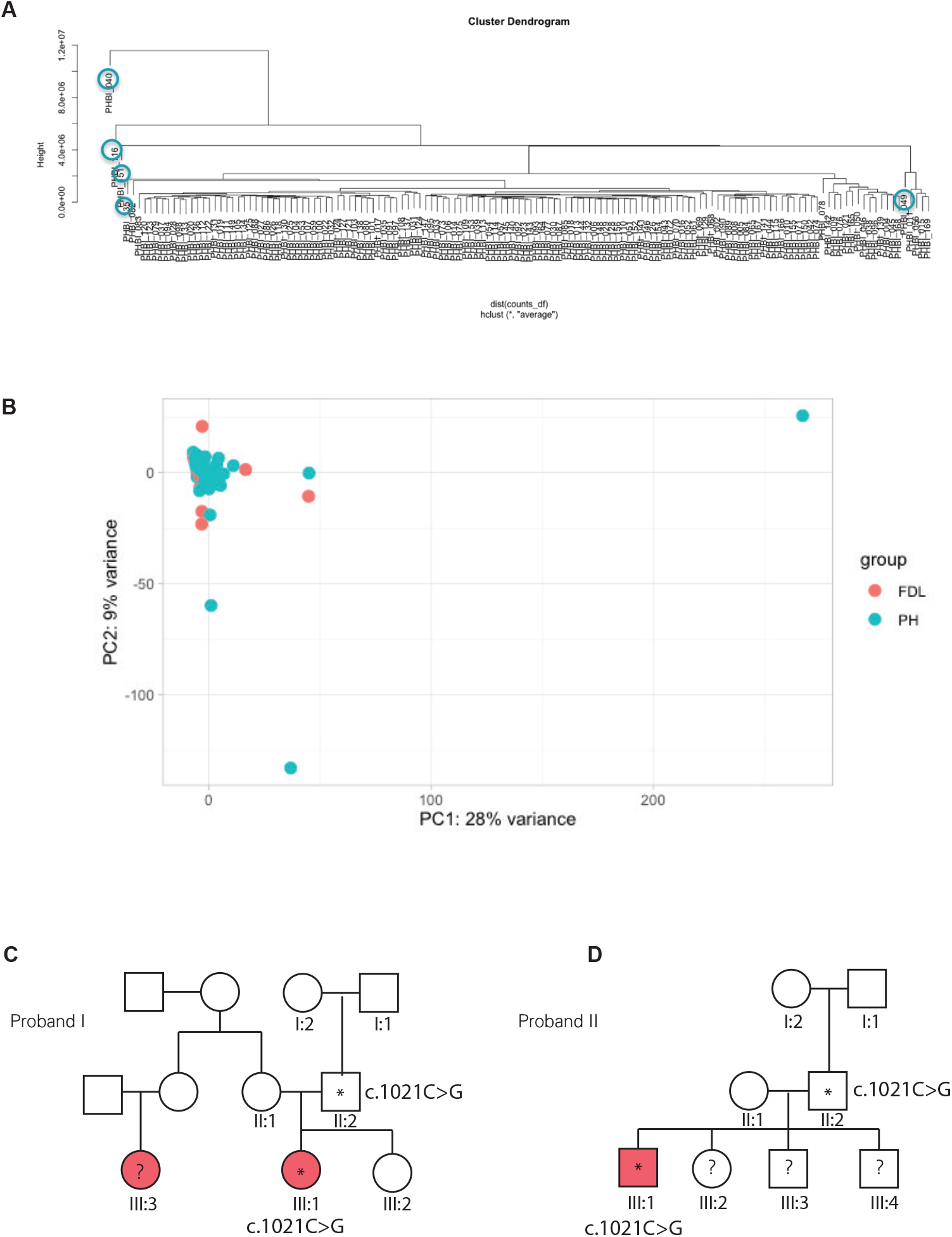
A-B) Cluster dendrogram (A) and principal component analysis (PCA) plot illustrating the outliers (circled in blue) in relation to the rest of the PHBI samples. FDL = Failed donor controls, PAH = Pulmonary arterial hypertension. C-D) Family pedigrees of index probands carrying FOXF1 variants. An asterisk represents a deleterious variant carrier; red color fill indicates the presence of a PAH syndrome; question marks represent undetermined genotypes.

**Supp. Figure 2:**
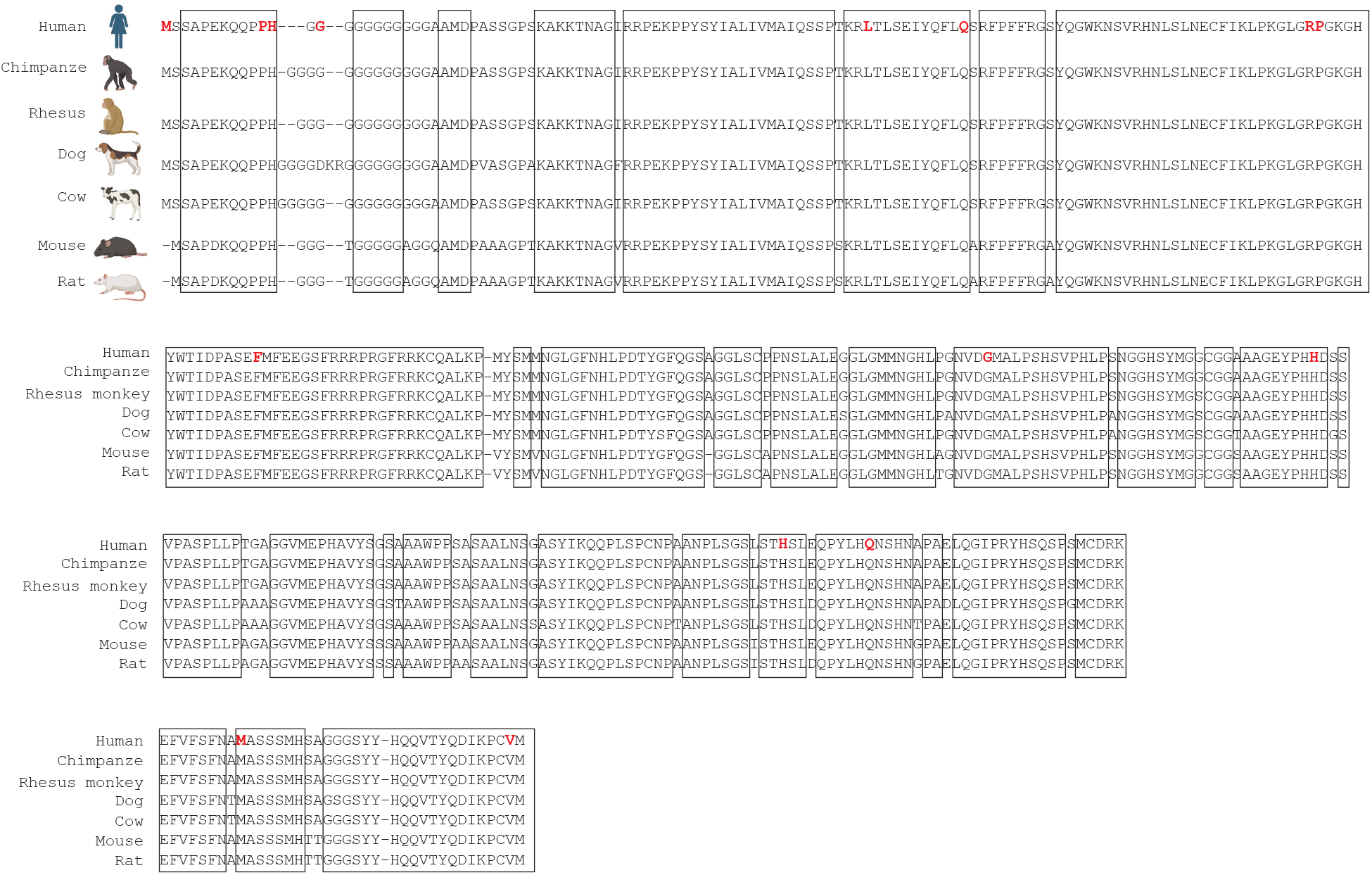
FOXF1 amino acid sequence across different eutherian species. Blocks represent areas of conservation across all species shown. Red residues indicate mutations in humans with PAH.

**Supp. Figure 3:**
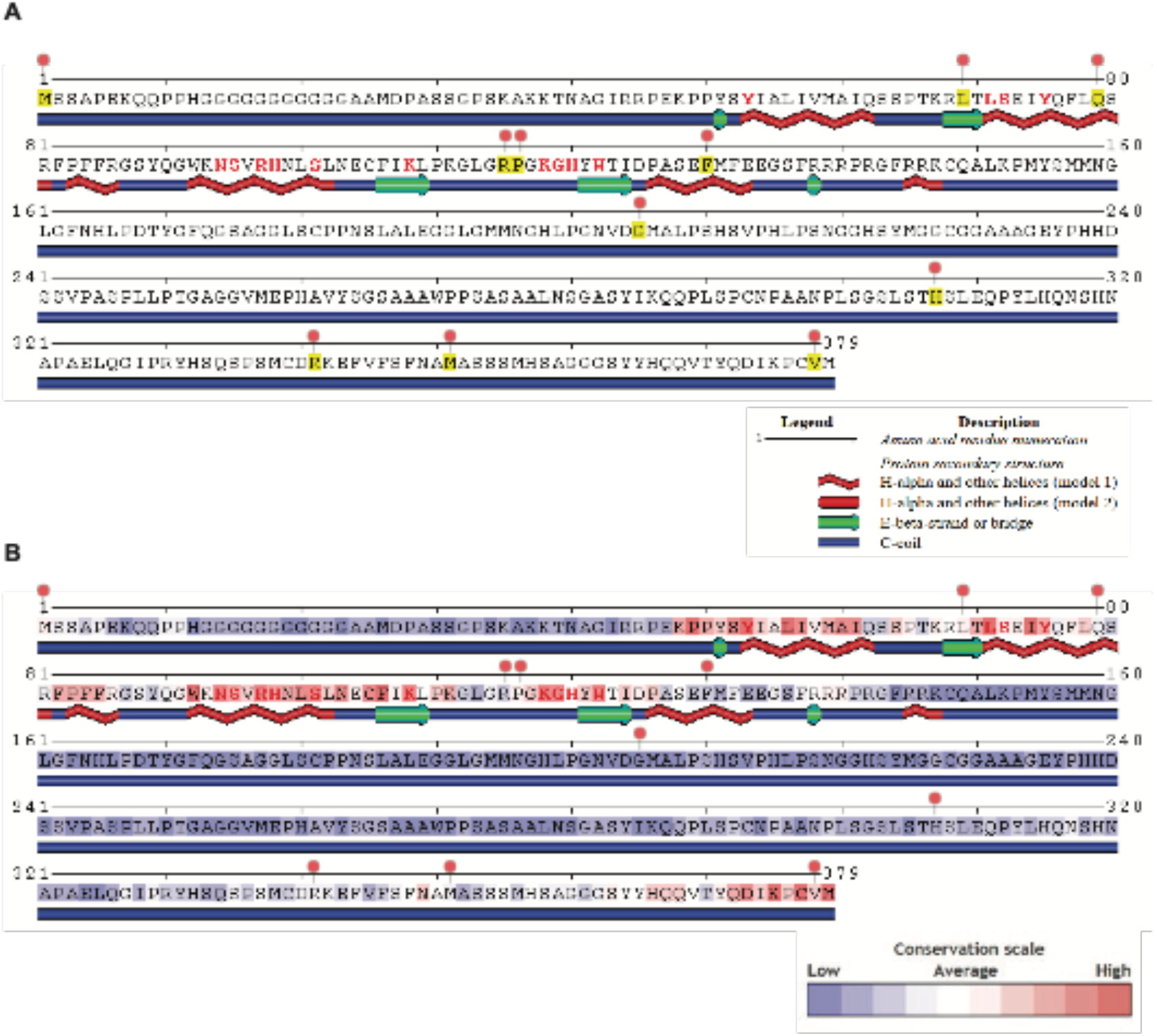
A) FOXF1 amino acid sequence with predicted secondary structures. Yellow residues (with lollipops) are mutated in patients with PAH. Red residues are predicted DNA binding sites. B) FOXF1 amino acid sequence showing conserved residues color-coded from low (blue) to high (red) on the conservation scale. Lollipops indicate mutated residues in patients with PAH.

**Supp. Figure 4:**
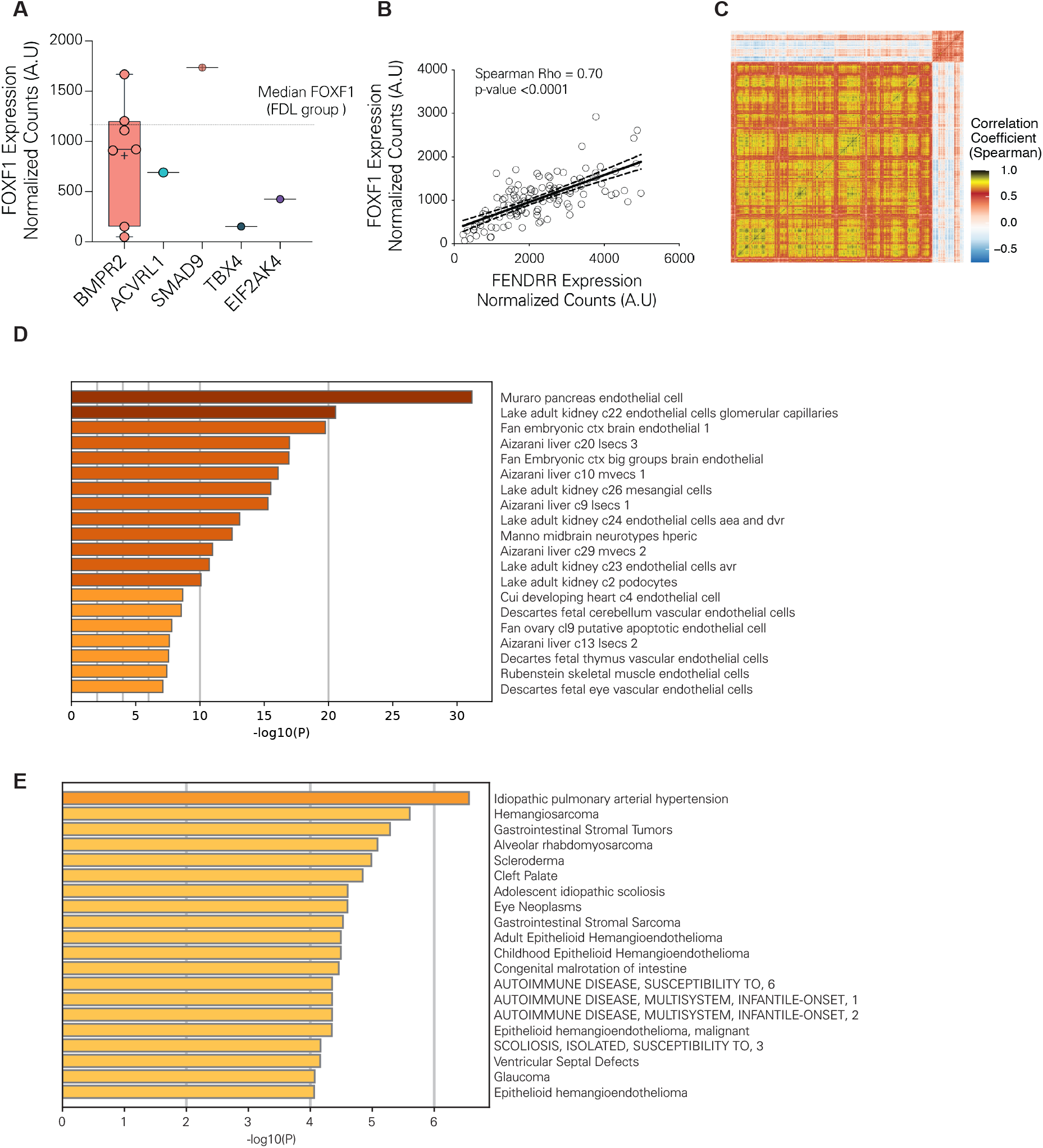
A) Box-and-whisker plots of FOXF1 normalized counts between PAH patients with known deleterious or pathogenic mutations in the selected genes. B) Correlation between FOXF1 and the long non-coding RNA FENDRR. A.U = Arbitrary units. C) Correlation matrix including all genes in the FOXF1 co-expression module. D) Bar chart of significantly enriched top cell signatures. The chart displays-log10(P) values for each dataset. E) Bar chart of significantly enriched DisGeNET associations. The chart displays-log10(P) values for each disease state.

**Supp. Figure 5:**
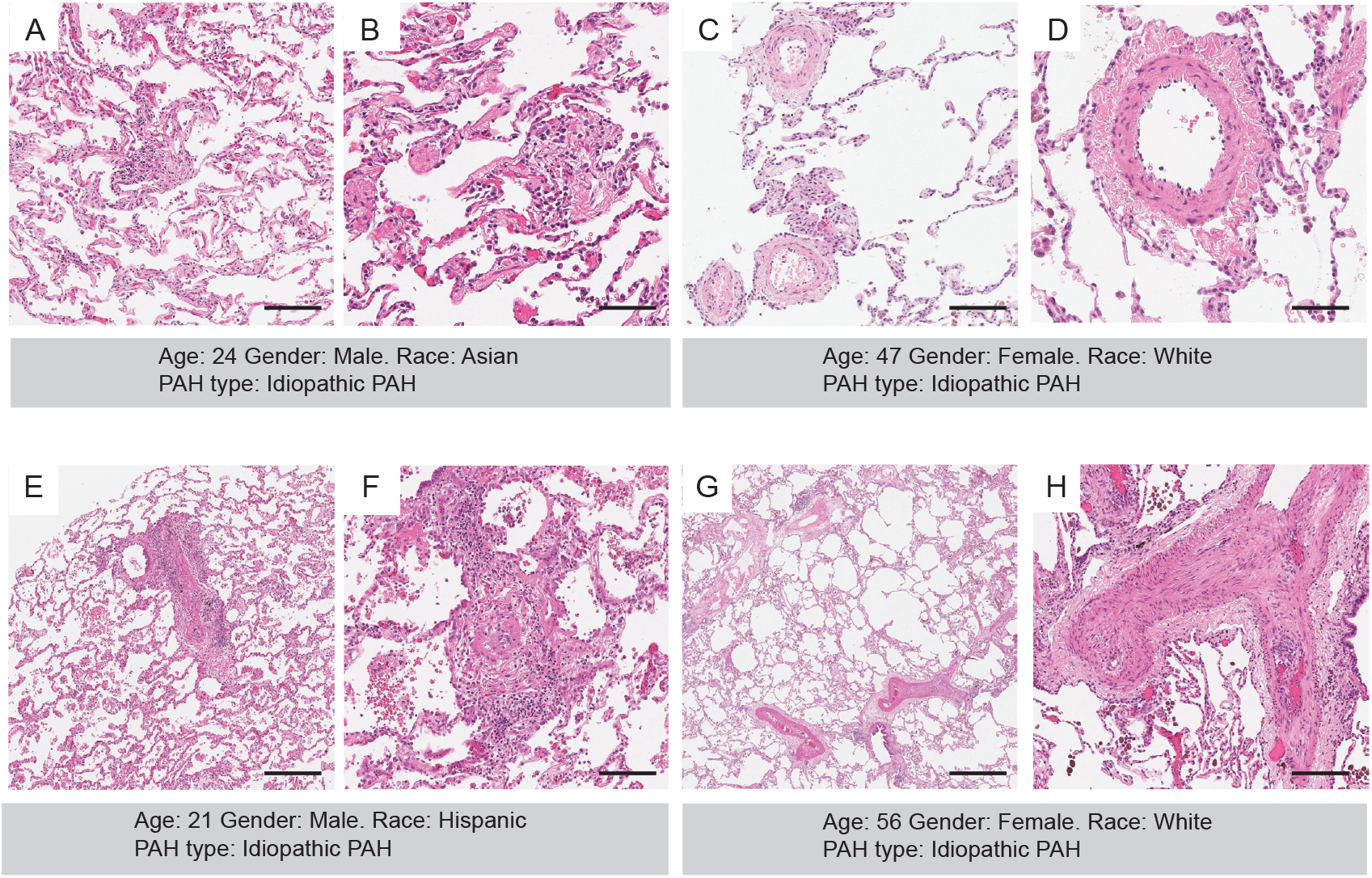
A-H) Hematoxylin and eosin stains of different PAH tissue samples showing various types of vascular remodeling consistent with PAH.

## References

1. Hassoun PM. Pulmonary Arterial Hypertension. N Engl J Med 2021;doi:10.1056/nejmra2000348.

2. Chang KY, Duval S, Badesch DB, Bull TM, Chakinala MM, Marco TD, et al. Mortality in Pulmonary Arterial Hypertension in the Modern Era: Early Insights From the Pulmonary Hypertension Association Registry. J Am Heart Assoc 2021;11:e024969.

3. Voelkel NF, Gomez-Arroyo J, Abbate A, Bogaard HJ, Nicolls MR. Pathobiology of pulmonary arterial hypertension and right ventricular failure. European Respiratory Journal 2012;40:1555–1565.

4. Hemnes AR, Humbert M. Pathobiology of pulmonary arterial hypertension: understanding the roads less travelled. Eur Respir Rev 2017;26:.

5. Nicolls MR, Voelkel NF. The Roles of Immunity in the Prevention and Evolution of Pulmonary Arterial Hypertension. A Perspective. Am J Respir Crit Care Med 2016;rccm.201608-1630PP.doi:10.1164/rccm.201608-1630pp.

6. Humbert M, Guignabert C, Bonnet S, Dorfmuller P, Klinger JR, Nicolls MR, et al. Pathology and pathobiology of pulmonary hypertension: state of the art and research perspectives. European Respiratory Journal 2019;53:.

7. Kwapiszewska G, Johansen AKZ, Gomez-Arroyo J, Voelkel NF. Role of the Aryl Hydrocarbon Receptor/ARNT/Cytochrome P450 System in Pulmonary Vascular Diseases. Circ Res 2019;125:356– 366.

8. Weir EK, Reeve HL, Huang JM, Michelakis E, Nelson DP, Hampl V, et al. Anorexic agents aminorex, fenfluramine, and dexfenfluramine inhibit potassium current in rat pulmonary vascular smooth muscle and cause pulmonary vasoconstriction. Circulation 1996;94:2216–2220.

9. Orcholski ME, Yuan K, Rajasingh C, Tsai H, Shamskhou EA, Dhillon NK, et al. Drug-induced pulmonary arterial hypertension: a primer for clinicians and scientists. AJP: Lung Cellular and Molecular Physiology 2018;314:L967–L983.

10. Dalvi P, Wang K, Mermis J, Zeng R, Sanderson M, Johnson S, et al. HIV-1/Cocaine Induced Oxidative Stress Disrupts Tight Junction Protein-1 in Human Pulmonary Microvascular Endothelial Cells: Role of Ras/ERK1/2 Pathway. 2014;

11. Montani D, Bergot E, Günther S, Savale L, Bergeron A, Bourdin A, et al. Pulmonary Arterial Hypertension in Patients Treated by Dasatinib. Circulation 2012;doi:10.1161/circulationaha.111.079921.

12. Marecki JC, Cool CD, Parr JE, Beckey VE, Luciw PA, Tarantal AF, et al. HIV-1 Nef is associated with complex pulmonary vascular lesions in SHIV-nef-infected macaques. Am J Respir Crit Care Med 2006;174:437–445.

13. Becker MO, Kill A, Kutsche M, Guenther J, Rose A, Tabeling C, et al. Vascular Receptor Autoantibodies in Pulmonary Arterial Hypertension Associated with Systemic Sclerosis. Am J Resp Crit Care 2014;190:808–817.

14. Humbert M, Monti G, Brenot F, Sitbon O, Portier A, Grangeot-Keros L, et al. Increased interleukin-1 and interleukin-6 serum concentrations in severe primary pulmonary hypertension. Am J Respir Crit Care Med 1995;151:1628–1631.

15. Shao D, Perros F, Caramori G, Meng C, Dormuller P, Chou P-C, et al. Nuclear IL-33 regulates soluble ST2 receptor and IL-6 expression in primary human arterial endothelial cells and is decreased in idiopathic pulmonary arterial hypertension. Biochem Biophys Res Commun 2014;451:8–14.

16. Heresi GA, Aytekin M, Hammel JP, Wang S, Chatterjee S, Dweik RA. Plasma interleukin-6 adds prognostic information in pulmonary arterial hypertension. European Respiratory Journal 2014;43:912–914.

17. Soon E, Crosby A, Southwood M, Yang P, Tajsic T, Toshner M, et al. BMPR-II Deficiency Promotes Pulmonary Hypertension via Increased Inflammatory Cytokine Production. Am J Respir Crit Care Med 2015;doi:10.1164/rccm.201408-1509oc.

18. Taraseviciene-Stewart L, Kasahara Y, Alger L, Hirth P, Mahon GM, Waltenberger J, et al. Inhibition of the VEGF receptor 2 combined with chronic hypoxia causes cell death-dependent pulmonary endothelial cell proliferation and severe pulmonary hypertension. FASEB J 2001;15:427–438.

19. Morrell NW, Aldred MA, Chung WK, Elliott CG, Nichols WC, Soubrier F, et al. Genetics and genomics of pulmonary arterial hypertension. European Respiratory Journal 2019;53:1801899.

20. Gräf S, Haimel M, Bleda M, Hadinnapola C, Southgate L, Li W, et al. Identification of rare sequence variation underlying heritable pulmonary arterial hypertension. Nat Commun 2018;9:.

21. Zhu N, Gonzaga-Jauregui C, Welch CL, Ma L, Qi H, King AK, et al. Exome Sequencing in Children With Pulmonary Arterial Hypertension Demonstrates Differences Compared With Adults. Circulation Genom Precis Medicine 2018;11:e001887.

22. Lévy M, Eyries M, Szezepanski I, Ladouceur M, Nadaud S, Bonnet D, et al. Genetic analyses in a cohort of children with pulmonary hypertension. European Respiratory Journal 2016;48:1118–1126.

23. Zhu N, Pauciulo MW, Welch CL, Lutz KA, Coleman AW, Gonzaga-Jauregui C, et al. Novel risk genes and mechanisms implicated by exome sequencing of 2572 individuals with pulmonary arterial hypertension. 2019;

24. Montani D, Lechartier B, Girerd B, Eyries M, Ghigna M-R, Savale L, et al. An emerging phenotype of pulmonary arterial hypertension patients carrying SOX17 variants. Eur Respir J 2022;60:2200656.

25. Postma AV, Rapp CK, Knoflach K, Volk AE, Lemke JR, Ackermann M, et al. Biallelic variants in the calpain regulatory subunit CAPNS1 cause pulmonary arterial hypertension. Genet Med Open 2023;1:100811.

26. Kalinichenko VV, Lim L, Stolz DB, Shin B, Rausa FM, Clark J, et al. Defects in Pulmonary Vasculature and Perinatal Lung Hemorrhage in Mice Heterozygous Null for the Forkhead Box f1 Transcription Factor. Dev Biol 2001;235:489–506.

27. Carlsson P, Mahlapuu M. Forkhead transcription factors: key players in development and metabolism. Dev Biol 2002;250:1–23.

28. Ren X, Ustiyan V, Pradhan A, Cai Y, Havrilak JA, Bolte CS, et al. FOXF1 Transcription Factor Is Required for Formation of Embryonic Vasculature by Regulating VEGF Signaling in Endothelial Cells. Circ Res 2014;doi:10.1161/circresaha.115.304382.

29. Sen P, Yang Y, Navarro C, Silva I, Szafranski P, Kolodziejska KE, et al. Novel FOXF1 mutations in sporadic and familial cases of alveolar capillary dysplasia with misaligned pulmonary veins imply a role for its DNA binding domain. Hum Mutat 2013;34:801–811.

30. Stankiewicz P, Sen P, Bhatt SS, Storer M, Xia Z, Bejjani BA, et al. Genomic and genic deletions of the FOX gene cluster on 16q24.1 and inactivating mutations of FOXF1 cause alveolar capillary dysplasia and other malformations. Am J Hum Genet 2009;84:780– 791.

31. Zhu N, Swietlik EM, Welch CL, Pauciulo MW, Hagen JJ, Zhou X, et al. Correction to: Rare variant analysis of 4241 pulmonary arterial hypertension cases from an international consortium implicates FBLN2, PDGFD, and rare de novo variants in PAH. Genome Med 2021;13:106.

32. Stacher E, Graham BB, Hunt JM, Gandjeva A, Groshong SD, McLaughlin VV, et al. Modern age pathology of pulmonary arterial hypertension. Am J Respir Crit Care Med 2012;186:261–272.

33. Symersky P, Jansen SMA, Kamminga SK, Meijboom LJ, Lust EJ, Eghtesady P, et al. Improvement in exercise capacity after a modified Potts shunt in an adult patient with pulmonary arterial hypertension. Erj Open Res 2021;7:00287–02021.

34. Karczewski KJ, Francioli LC, Tiao G, Cummings BB, Alföldi J, Wang Q, et al. The mutational constraint spectrum quantified from variation in 141,456 humans. Nature 2020;581:434–443.

35. Kircher M, Witten DM, Jain P, O’Roak BJ, Cooper GM, Shendure J. A general framework for estimating the relative pathogenicity of human genetic variants. Nat Genet 2014;46:310– 315.

36. Rentzsch P, Witten D, Cooper GM, Shendure J, Kircher M. CADD: predicting the deleteriousness of variants throughout the human genome. Nucleic Acids Res 2019;47:D886–D894.

37. Lambert SA, Jolma A, Campitelli LF, Das PK, Yin Y, Albu M, et al. The Human Transcription Factors. Cell 2018;172:650–665.

38. Havrilla JM, Pedersen BS, Layer RM, Quinlan AR. A map of constrained coding regions in the human genome. Nat Genet 2019;51:88–95.

39. Dharmadhikari AV, Szafranski P, Kalinichenko VV, Stankiewicz P. Genomic and Epigenetic Complexity of the FOXF1 Locus in 16q24.1: Implications for Development and Disease. Curr Genomics 2015;16:107–116.

40. Dai S, Qu L, Li J, Chen Y. Toward a mechanistic understanding of DNA binding by forkhead transcription factors and its perturbation by pathogenic mutations. Nucleic Acids Res 2021;49:10235–10249.

41. Hong J, Wong B, Rhodes CJ, Kurt Z, Schwantes-An T-H, Mickler EA, et al. Integrative Multiomics to Dissect the Lung Transcriptional Landscape of Pulmonary Arterial Hypertension. bioRxiv 2023;2023.01.12.523812.doi:10.1101/2023.01.12.523812.

42. Stearman RS, Bui QM, Speyer G, Handen A, Cornelius AR, Graham BB, et al. Systems Analysis of the Human Pulmonary Arterial Hypertension Lung Transcriptome. Am J Resp Cell Mol 2018;60:637–649.

43. Szafranski P, Gambin T, Karolak JA, Popek E, Stankiewicz P. Lung-specific distant enhancer cis regulates expression of FOXF1 and lncRNA FENDRR. Hum Mutat 2021;42:694–698.

44. Slot E, Edel G, Cutz E, Heijst Avan, Post M, Schnater M, et al. Alveolar capillary dysplasia with misalignment of the pulmonary veins: clinical, histological, and genetic aspects. Pulmonary Circulation 2018;8:204589401879514.

45. RR N, D P, Z Y, M E. Congenital Alveolar Capillary Dysplasia and New Associations: A Case Series with a Report of New Associations and Literature Review. Méd Rep Case Stud 2017;2:1– 6.

46. Onda T, Manabe A, Akimoto T, Hayasaka I, Ikeda M, Furuse Y, et al. Incidence of alveolar capillary dysplasia with misalignment of pulmonary veins in infants with unexplained severe pulmonary hypertension: The roles of clinical, pathological, and genetic testing. Early Hum Dev 2021;155:105323.

47. Towe CT, White FV, Grady RM, Sweet SC, Eghtesady P, Wegner DJ, et al. Infants with Atypical Presentations of Alveolar Capillary Dysplasia with Misalignment of the Pulmonary Veins Who Underwent Bilateral Lung Transplantation. J Pediatr 2018;194:158-164.e1.

48. Reiter J, Szafranski P, Breuer O, Perles Z, Dagan T, Stankiewicz P, et al. Variable phenotypic presentation of a novel FOXF1 missense mutation in a single family. Pediatr Pulmonol 2016;51:921–927.

49. Kalinichenko VV, Gusarova GA, Kim I-M, Shin B, Yoder HM, Clark J, et al. Foxf1 haploinsufficiency reduces Notch-2 signaling during mouse lung development. Am J Physiol-Lung Cell Mol Physiol 2004;286:L521–L530.

50. Pradhan A, Dunn A, Ustiyan V, Bolte C, Wang G, Whitsett JA, et al. The S52F FOXF1 Mutation Inhibits STAT3 Signaling and Causes Alveolar Capillary Dysplasia. Am J Respir Crit Care Med 2019;rccm.201810-1897OC.doi:10.1164/rccm.201810-1897oc.

51. Langfelder P, Horvath S. WGCNA: an R package for weighted correlation network analysis. BMC Bioinform 2008;9:559.

52. Guo M, Wikenheiser-Brokamp KA, Kitzmiller JA, Jiang C, Wang G, Wang A, et al. Single Cell Multiomics Identifies Cells and Genetic Networks Underlying Alveolar Capillary Dysplasia. Am J Respir Crit Care Med 2023;doi:10.1164/rccm.202210-2015oc.

53. Guo M, Wikenheiser-Brokamp KA, Kitzmiller JA, Jiang C, Wang G, Wang A, et al. Single Cell Multiomics Identifies Cells and Genetic Networks Underlying Alveolar Capillary Dysplasia. Am J Respir Crit Care Med 2023;208:709–725.

54. Sikkema L, Ramírez-Suástegui C, Strobl DC, Gillett TE, Zappia L, Madissoon E, et al. An integrated cell atlas of the lung in health and disease. Nat Med 2023;29:1563–1577.

55. Gillich A, Zhang F, Farmer CG, Travaglini KJ, Tan SY, Gu M, et al. Capillary cell-type specialization in the alveolus. Nature 2020;586:785–789.

56. Zhou Y, Zhou B, Pache L, Chang M, Khodabakhshi AH, Tanaseichuk O, et al. Metascape provides a biologist-oriented resource for the analysis of systems-level datasets. Nat Commun 2019;10:1523.

57. Piñero J, Bravo À, Queralt-Rosinach N, Gutiérrez-Sacristán A, Deu-Pons J, Centeno E, et al. DisGeNET: a comprehensive platform integrating information on human disease-associated genes and variants. Nucleic Acids Res 2017;45:D833–D839.

58. Ardini-Poleske ME, Clark RF, Ansong C, Carson JP, Corley RA, Deutsch GH, et al. LungMAP: The Molecular Atlas of Lung Development Program. Am J Physiol-Lung Cell Mol Physiol 2017;313:L733–L740.

59. Dharmadhikari AV, Sun JJ, Gogolewski K, Carofino BL, Ustiyan V, Hill M, et al. Lethal lung hypoplasia and vascular defects in mice with conditional Foxf1 overexpression. Biol Open 2016;5:1595–1606.

60. Nassar LR, Barber GP, Benet-Pagès A, Casper J, Clawson H, Diekhans M, et al. The UCSC Genome Browser database: 2023 update. Nucleic Acids Res 2022;51:D1188–D1195.

61. Rauluseviciute I, Riudavets-Puig R, Blanc-Mathieu R, Castro-Mondragon JA, Ferenc K, Kumar V, et al. JASPAR 2024: 20th anniversary of the open-access database of transcription factor binding profiles. Nucleic Acids Res 2023;52:D174–D182.

62. Matys V. TRANSFAC(R) and its module TRANSCompel(R): transcriptional gene regulation in eukaryotes. Nucleic Acids Res 2006;34:D108–D110.

63. Humbert M, McLaughlin V, Gibbs JSR, Gomberg-Maitland M, Hoeper MM, Preston IR, et al. Sotatercept for the Treatment of Pulmonary Arterial Hypertension. N Engl J Med 2021;384:1204– 1215.

64. Yung LM, Yang P, Joshi S, Augur ZM, Kim SSJ, Bocobo GA, et al. ACTRIIA-Fc rebalances activin/GDF versus BMP signaling in pulmonary hypertension. Sci Transl Med 2020;12:.

65. Herman L, Todeschini A-L, Veitia RA. Forkhead Transcription Factors in Health and Disease. Trends Genet 2021;37:460–475.

66. Jackson BC, Carpenter C, Nebert DW, Vasiliou V. Update of human and mouse forkhead box (FOX) gene families. Human genomics 2010;4:345–352.

67. Cai Y, Bolte C, Le T, Goda C, Xu Y, Kalin TV, et al. FOXF1 maintains endothelial barrier function and prevents edema after lung injury. Sci Signal 2016;9:ra40.

68. Wang G, Wen B, Deng Z, Zhang Y, Kolesnichenko OA, Ustiyan V, et al. Endothelial progenitor cells stimulate neonatal lung angiogenesis through FOXF1-mediated activation of BMP9/ACVRL1 signaling. Nat Commun 2022;13:2080.

69. Trembath RC, Thomson JR, Machado RD, Morgan NV, Atkinson C, Winship I, et al. Clinical and molecular genetic features of pulmonary hypertension in patients with hereditary hemorrhagic telangiectasia. N Engl J Med 2001;345:325–334.

70. Grynblat J, Bogaard H-J, Eyries M, Meyrignac O, Savale L, Jais X, et al. Pulmonary vascular phenotype identified in patients with GDF2 (BMP9) or BMP10 variants: An international multicentre study. Eur Respir J 2024;2301634.doi:10.1183/13993003.01634-2023.

71. Hodgson J, Swietlik EM, Salmon RM, Hadinnapola C, Nikolic I, Wharton J, et al. Characterization of GDF2 Mutations and Levels of BMP9 and BMP10 in Pulmonary Arterial Hypertension. Am J Resp Crit Care 2020;201:575–585.

72. Isobe S, Nair RV, Kang HY, Wang L, Moonen J-R, Shinohara T, et al. Reduced FOXF1 links unrepaired DNA damage to pulmonary arterial hypertension. Nat Commun 2023;14:7578.

73. Bishop NB, Stankiewicz P, Steinhorn RH. Alveolar capillary dysplasia. Am J Respir Crit Care Med 2011;184:172–179.

74. Sirianansopa K, Prasertsan P, Ruangnapa K, Saelim K, Koranantakul P. Unusual presentation of alveolar capillary dysplasia with misalignment of the pulmonary veins in a child with respiratory syncytial virus pneumonia: A case report. Respirol Case Rep 2023;11:e01089.

75. Abdallah HI, Karmazin N, Marks LA. Late Presentation of Misalignment of Lung Vessels with Alveolar Capillary Dysplasia. Critical Care Medicine 1993;21:628–630.

76. Chelliah BP, Brown D, Cohen M, Talleyrand AJ, Shen-Schwarz S. Alveolar capillary dysplasia--a cause of persistent pulmonary hypertension unresponsive to a second course of extracorporeal membrane oxygenation. Pediatrics 1995;96:1159–1161.

77. Shankar V, Haque A, Johnson J, Pietsch J. Late presentation of alveolar capillary dysplasia in an infant. Pediatr Crit Care Med 2006;7:177–179.

78. Mahlapuu M, Pelto-Huikko M, Aitola M, Enerbäck S, Carlsson P. FREAC-1 Contains a Cell-Type-Specific Transcriptional Activation Domain and Is Expressed in Epithelial–Mesenchymal Interfaces. Dev Biol 1998;202:183–195.

79. Bölükbaşı EY, Karolak JA, Szafranski P, Gambin T, Matsika A, McManus S, et al. Variable expressivity in a four-generation ACDMPV family with a non-coding hypermorphic SNV in trans to the frameshifting FOXF1 variant. Eur J Hum Genet 2022;1– 5.doi:10.1038/s41431-022-01159-x.

80. Sen P, Gerychova R, Janku P, Jezova M, Valaskova I, Navarro C, et al. A familial case of alveolar capillary dysplasia with misalignment of pulmonary veins supports paternal imprinting of FOXF1 in human. Eur J Hum Genet 2013;21:474–477.

81. Szafranski P, Liu Q, Karolak JA, Song X, Leeuw Nde, Faas B, et al. Association of rare non-coding SNVs in the lung-specific FOXF1 enhancer with a mitigation of the lethal ACDMPV phenotype. Hum Genet 2019;138:1301–1311.

82. Szafranski P, Gambin T, Dharmadhikari AV, Akdemir KC, Jhangiani SN, Schuette J, et al. Pathogenetics of alveolar capillary dysplasia with misalignment of pulmonary veins. Hum Genet 2016;135:569–586.

83. Eyries M, Montani D, Girerd B, Favrolt N, Riou M, Faivre L, et al. Familial pulmonary arterial hypertension by KDR heterozygous loss of function. European Respiratory Journal 2020;55:.

84. Haarman MG, Kerstjens-Frederikse WS, Berger RMF. TBX4 variants and pulmonary diseases: getting out of the ‘Box.’ Curr Opin Pulm Med 2020;26:277–284.

85. Maddaloni C, Ronci S, Rose DUD, Bersani I, Campi F, Nardo MD, et al. Neonatal persistent pulmonary hypertension related to a novel TBX4 mutation: case report and review of the literature. Ital J Pediatr 2024;50:41.

86. Galambos C, Mullen MP, Shieh JT, Schwerk N, Kielt MJ, Ullmann N, et al. Phenotype characterisation of TBX4 mutation and deletion carriers with neonatal and paediatric pulmonary hypertension. Eur Respir J 2019;54:1801965.

87. Tamura M, Sasaki Y, Koyama R, Takeda K, Idogawa M, Tokino T. Forkhead transcription factor FOXF1 is a novel target gene of the p53 family and regulates cancer cell migration and invasiveness. Oncogene 2014;33:4837–4846.

88. Sun F, Wang G, Pradhan A, Xu K, Gomez-Arroyo J, Zhang Y, et al. Nanoparticle Delivery of STAT3 Alleviates Pulmonary Hypertension in a Mouse Model of Alveolar Capillary Dysplasia. Circulation 2021;144:539–555.

89. Long L, Ormiston ML, Yang X, Southwood M, Gräf S, Machado RD, et al. Selective enhancement of endothelial BMPR-II with BMP9 reverses pulmonary arterial hypertension. Nat Med 2015;doi:10.1038/nm.3877.

90. Rai PR, Cool CD, King JAC, Stevens T, Burns N, Winn RA, et al. The cancer paradigm of severe pulmonary arterial hypertension. Am J Respir Crit Care Med 2008;178:558–564.

91. Lo P-K, Lee JS, Sukumar S. The p53–p21WAF1 checkpoint pathway plays a protective role in preventing DNA rereplication induced by abrogation of FOXF1 function. Cell Signal 2012;24:316–324.

92. Lo P-K, Lee JS, Liang X, Han L, Mori T, Fackler MJ, et al. Epigenetic Inactivation of the Potential Tumor Suppressor Gene FOXF1 in Breast Cancer. Cancer Res 2010;70:6047–6058.

93. Swietlik EM, Greene D, Zhu N, Megy K, Cogliano M, Rajaram S, et al. Bayesian Inference Associates Rare KDRVariants with Specific Phenotypes in Pulmonary Arterial Hypertension. Circ: Genomic and Precision Medicine 2020;144:275.

94. Rhodes CJ, Batai K, Bleda M, Haimel M, Southgate L, Germain M, et al. Genetic determinants of risk in pulmonary arterial hypertension: international genome-wide association studies and meta-analysis. Lancet Respir Med 2019;7:227–238.

95. Austin ED, Ma L, LeDuc C, Rosenzweig EB, Borczuk A, III Jap, et al. Whole Exome Sequencing to Identify a Novel Gene (Caveolin-1) Associated With Human Pulmonary Arterial Hypertension. Circulation Cardiovasc Genetics 2012;5:336–343.

96. Ma L, Roman-Campos D, Austin ED, Eyries M, Sampson KS, Soubrier F, et al. A novel channelopathy in pulmonary arterial hypertension. N Engl J Med 2013;369:351–361.

97. Li H, Durbin R. Fast and accurate short read alignment with Burrows–Wheeler transform. Bioinformatics 2009;25:1754–1760.

98. McKenna A, Hanna M, Banks E, Sivachenko A, Cibulskis K, Kernytsky A, et al. The Genome Analysis Toolkit: A MapReduce framework for analyzing next-generation DNA sequencing data. Genome Res 2010;20:1297–1303.

99. Lek M, Karczewski KJ, Minikel EV, Samocha KE, Banks E, Fennell T, et al. Analysis of protein-coding genetic variation in 60,706 humans. Nature 2016;536:285–291.

100. Landrum MJ, Lee JM, Benson M, Brown GR, Chao C, Chitipiralla S, et al. ClinVar: improving access to variant interpretations and supporting evidence. Nucleic Acids Res 2017;46:gkx1153..

101. Liu X, Wu C, Li C, Boerwinkle E. dbNSFP v3.0: A One-Stop Database of Functional Predictions and Annotations for Human Nonsynonymous and Splice-Site SNVs. Hum Mutat 2016;37:235– 241.

102. Li C, Zhi D, Wang K, Liu X. MetaRNN: differentiating rare pathogenic and rare benign missense SNVs and InDels using deep learning. Genome Med 2022;14:115.

103. Jagadeesh KA, Wenger AM, Berger MJ, Guturu H, Stenson PD, Cooper DN, et al. M-CAP eliminates a majority of variants of uncertain significance in clinical exomes at high sensitivity. Nat Genet 2016;48:1581–1586.

104. Cheng J, Novati G, Pan J, Bycroft C, Žemgulytė A, Applebaum T, et al. Accurate proteome-wide missense variant effect prediction with AlphaMissense. Science 2023;381:eadg7492.

105. Kopanos C, Tsiolkas V, Kouris A, Chapple CE, Aguilera MA, Meyer R, et al. VarSome: the human genomic variant search engine. Bioinformatics 2019;35:1978–1980.

106. MacArthur DG, Manolio TA, Dimmock DP, Rehm HL, Shendure J, Abecasis GR, et al. Guidelines for investigating causality of sequence variants in human disease. Nature 2014;508:469–476.

107. Richards S, Aziz N, Bale S, Bick D, Das S, Gastier-Foster J, et al. Standards and guidelines for the interpretation of sequence variants: a joint consensus recommendation of the American College of Medical Genetics and Genomics and the Association for Molecular Pathology. Genet Med Genet Med; 2015. p. 405–424.

108. Germain M, Eyries M, Montani D, Poirier O, Girerd B, Dorfmuller P, et al. Genome-wide association analysis identifies a susceptibility locus for pulmonary arterial hypertension. Nat Genet 2013;45:518–521.

109. Machado RD, Eickelberg O, Elliott CG, Geraci MW, Hanaoka M, Loyd JE, et al. Genetics and genomics of pulmonary arterial hypertension. J Am Coll Cardiol 2009;54:S32–42.

110. Zhu N, Welch CL, Wang J, Allen PM, Gonzaga-Jauregui C, Ma L, et al. Rare variants in SOX17 are associated with pulmonary arterial hypertension with congenital heart disease. Genome Med 2018;10:56.

111. Varadi M, Anyango S, Deshpande M, Nair S, Natassia C, Yordanova G, et al. AlphaFold Protein Structure Database: massively expanding the structural coverage of protein-sequence space with high-accuracy models. Nucleic Acids Res 2021;50:D439–D444.

112. Porollo AA, Adamczak R, Meller J. POLYVIEW: a flexible visualization tool for structural and functional annotations of proteins. Bioinformatics 2004;20:2460–2462.

113. Bray NL, Pimentel H, Melsted P, Pachter L. Near-optimal probabilistic RNA-seq quantification. Nat Biotechnol 2016;34:525– 527.

114. Soneson C, Love MI, Robinson MD. Differential analyses for RNA-seq: transcript-level estimates improve gene-level inferences. F1000Res 2015;4:1521–19.

115. Love MI, Huber W, Anders S. Moderated estimation of fold change and dispersion for RNA-seq data with DESeq2. Genome Biol 2014;15:31.

116. Love MI, Anders S, Kim V, Huber W. RNA-Seq workflow: gene-level exploratory analysis and differential expression. F1000Res 2015;doi:10.12688/f1000research.7035.1.

117. Zheng R, Wan C, Mei S, Qin Q, Wu Q, Sun H, et al. Cistrome Data Browser: expanded datasets and new tools for gene regulatory analysis. Nucleic Acids Res 2019;47:D729–D735.

118. Weirauch MT, Yang A, Albu M, Cote AG, Montenegro-Montero A, Drewe P, et al. Determination and Inference of Eukaryotic Transcription Factor Sequence Specificity. Cell 2014;158:1431–1443.

